# Enduring Autism-like Phenotypes and Deregulated Hypothalamic Prosocial Peptides After Early-Life Exposure to Indoor Flame Retardants in Male C57BL/6 Mice

**DOI:** 10.1101/2025.09.02.673882

**Authors:** Elena V. Kozlova, Gwendolyn M. Gonzalez, Maximillian E. Denys, Anthony E. Bishay, Roberto Gutierrez, Jack Reid, Julia M. Krum, Gregory Lampel, Naran Luvsanravdan, Kayhon M. Rabbani, Jordan Tu, Luis Campoy, Laura M. Anchondo, Crystal N. Luna, Duraan S. Olomi, Eduardo Monarrez, Valeria Carrillo, Jasmin D. Tran, Damon Platt, Yash Korde, Bhuvaneswari D. Chinthirla, Maher Blaibel, Simon Kim, Gladys Chompre, Allison L. Phillips, Heather M. Stapleton, Bernhard Henkelmann, Karl-Werner Schramm, Margarita C. Curras-Collazo

**Author notes:** **Corresponding author:** Dr. Margarita C. Curras-Collazo, Ph.D, Professor of Neuroscience, Department Molecular, Cell and Systems Biology, University of California, Riverside, Riverside, CA 92521.

## Abstract

**Background:** Polybrominated diphenyl ethers (PBDEs) are neuroendocrine disrupting chemicals that produce adverse neurodevelopmental effects. PBDEs have been implicated as risk factors for autism spectrum disorder (ASD), which is characterized by abnormal psychosocial functioning and is commonly accompanied by co-morbidities such as cognitive and attentional deficits. Here, we used a mouse model with translationally relevant exposure to establish direct causal evidence that maternal transfer of a commercial mixture of PBDEs, DE-71, produces ASD-relevant behavioral and neurochemical deficits in male offspring.

**Methods:** C57Bl6/N mouse dams were exposed to a commercial PBDE mixture, DE-71, via oral administration of 0 (vehicle control, VEH/CON), 0.1 (L-DE-71), or 0.4 (H-DE-71) mg/kg bw/d for 10 weeks, spanning three weeks prior to gestation through the end of lactation at postnatal day (PND) 21.

**Results:** Mass spectrometric analysis indicated dose-dependent transfer of PBDEs (in ppb) to brains of F1 male offspring at PND 30, with reduction in levels by PND 110. Adult F1 male offspring displayed ASD-relevant neurobehavioral phenotypes, including impaired short- and long-term social recognition memory (SRM), despite intact general sociability, and exaggerated repetitive behavior. Exposed mice also displayed altered olfactory discrimination of social odors, impaired novel object recognition memory, and reduced open field habituation. However, no changes were observed in anxiety-like, sensorimotor, or depressive-like behaviors relative to VEH/CON. At the molecular level, DE-71 exposed males displayed deregulated gene markers of prosocial neuropeptides. *Oxt* was upregulated in the paraventricular nucleus (PVN); *Avp* was upregulated in the PVN and bed nucleus of the stria terminalis (BNST) but downregulated in the lateral septum (LS); *Avp1ar* and *Adcyap1* were upregulated in the BNST; and *Adcyap1r1* was upregulated in the PVN, supraoptic nucleus (SON), and BNST.

**Conclusions:** These findings demonstrate that developmental PBDE exposure produces enduring behavioral and neurochemical phenotypes that resemble core domains of ASD, which may result from early neurodevelopmental reprogramming within central social and memory networks.

## Introduction

Autism spectrum disorder (ASD) is a clinically heterogeneous neurodevelopmental disorder (NDD) diagnosed based on deficits across two core symptom domains: social communication and interaction, and restricted, repetitive patterns of behavior, interests, or activities (American Psychiatric Association, 2013; Lord et al., 2018). ASD is frequently accompanied by comorbidities such as intellectual disability, attention deficit hyperactivity disorder (ADHD), and olfactory impairments (Khachadourian et al., 2023; Larsson et al., 2017). According to the US Centers for Disease Control (CDC), the prevalence of ASD has increased fivefold since 2000 and currently affects 1 in 31 children, with a disproportionately male bias (male-to-female ratio 3.4:1) (Shaw et al., 2025). This epidemic-like rise inASD prevalence cannot be fully explained by changes in diagnostic criteria and improved detection and awareness (Hertz-Picciotto & Delwiche, 2009). While ASD has strong genetic underpinnings, genetic factors contribute an estimated 40-80% of ASD heritability (Rylaarsdam & Guemez-Gamboa, 2019). Moreover, findings from monozygotic twin studies show that genetics alone cannot account for ASD risk, suggesting a role for environmental factors (Isaksson et al., 2022). Thus, the etiology of ASD is hypothesized to involve gene–environment (G × E) interactions, with environmental exposures acting alongside genetic susceptibility (Keil-Stietz & Lein, 2023); however, the complex interactions between toxicants and genetic factors remain understudied, highlighting a critical gap in understanding ASD risk.

Mounting evidence shows significant associations between early-life exposure to endocrine-disrupting chemicals (EDCs) and NDD-risk (Braun, 2017; Herbstman & Mall, 2014). Polybrominated diphenyl ethers (PBDEs) are a class of persistent organic pollutants (POPs) used in a wide range of consumer products including building materials, electronics, textiles, plastics, and foams (Alaee et al., 2003). PBDEs have been detected in human serum, placenta, and breastmilk, and are readily transferred to the developing fetus, raising concern for their neurodevelopmental toxicity (Costa & Giordano, 2007). PBDEs have been banned or voluntarily phased out, leading to slow, but measurable, declines in environmental and biota concentrations of some congeners (Drage et al., 2019; Guo et al., 2016). However, PBDE contamination is predicted to remain an ongoing problem for decades to come due to their long half-lives, persistence in e-waste (Ohajinwa et al., 2019), and their inadvertent reappearance into the environment via recycling into consumer products (Abbasi et al., 2019).

Findings in humans and animals have raised concerns about PBDE exposure in the framework of the developmental origins of health and disease hypothesis, suggesting that PBDE exposure during early developmental windows of biological plasticity may disproportionately disrupt neurobehavioral outcomes (Barker, 1995; Heindel & Vandenberg, 2015). Epidemiological studies have found that developmental PBDE exposure is associated with poor executive function, lower IQ, attention difficulties, and disrupted behavioral regulation, reported as impulsivity and reduced social competence, suggesting that PBDEs may negatively impact children’s social development (Braun et al., 2014; Ding et al., 2015; Gascon et al., 2011; Gibson et al., 2018; Messer, 2010). In the HOME longitudinal cohort, serum levels of PBDEs in pregnant mothers (2003-2006) were associated with greater (BDE-28) or fewer (BDE-85) autistic behaviors in their toddlers (Braun et al., 2014). Findings from the HOME study also recently revealed that male adolescents whose mothers had an increased gestational serum ∑_5_BDE concentration exhibit decreased social competence (Hartley et al., 2022). Postnatal BDE-47 exposure was associated with a significantly higher risk of poor social competence symptoms in a cohort of 4 year olds from Spain (Gascon et al., 2011). Similarly, in the EARLI study, higher maternal BDE-47 was associated with reduced social reciprocity scores (SRS) (Song et al., 2023).

Experimental studies in murine models have provided supporting evidence that specific PBDE congeners adversely affect locomotor behavior, learning, and memory in offspring (Costa & Giordano, 2007), (Kodavanti & Curras-Collazo, 2010), (Pinson et al., 2016). However, data about the negative impact of PBDEs on psycho-social behavior remain limited. Broader investigations of brominated frame retardants (BFR) reported disrupted social cognition across several ASD-relevant domains (Gillera et al., 2020; Kim et al., 2015; Witchey et al., 2020), although other studies found no significant effects. These inconsistencies may be explained by the absence of a second ‘hit’, the BFR type, timing of exposure, sex and/or model (Gillera et al., 2020; Z. Li et al., 2021; Witchey et al., 2020; Woods et al., 2012). We recently reported that PBDE-exposed F1 female progeny display ASD-relevant behavioral characteristics concomitant with deregulated markers for prosocial hypothalamic genes (Kozlova et al., 2021), however, whether exposure in male offspring produces similar effects has not been studied.

Oxytocin (OXT) and vasopressin (AVP) are neuropeptides produced in the paraventricular (PVN) and supraoptic (SON) nuclei of the hypothalamic neurohypophyseal system (HNS) where they play essential roles across multiple social cognition domains such as social memory, social/emotional recognition and social reward (Ferguson et al., 2001); (Raam et al., 2017); (Bielsky et al., 2004); (Ferretti et al., 2019). Both neuropeptides can be altered in autistic individuals (Cataldo et al., 2018; László et al., 2023; Meyer-Lindenberg et al., 2011) and also used in the treatment of psychosocial disorders (Hendaus et al., 2019; Shahrestani et al., 2013). Evidence from animal models shows that thyroid hormones (THs) regulate hypothalamic OXT/AVP mRNA and content (Adan et al., 1992; Vasudevan et al., 2001), and axonal release (Adan et al., 1992; Ciosek & Drobnik, 2004; Dellovade et al., 1999; Vasudevan et al., 2001). We and others have also shown that the rat and the mouse neuroendocrine PVN and SON are vulnerable to disruption by thyroid-hormone-disrupting PBDEs and other EDCs, in parallel with social behavior deficits (Coburn et al., 2005, 2007; Garduño-Gutiérrez et al., 2023; Kozlova et al., 2021; Mucio-Ramírez et al., 2017; Patisaul, 2017; Reilly et al., 2022). PBDEs, along with structurally related polychlorinated biphenyls (PCBs), such as those found in Aroclor 1254, reduce physiologically-stimulated AVP release from the somatodendritic compartments of SON magnocellular neuroendocrine neurons (Coburn et al., 2005, 2007). This suggests that PBDEs may contribute to ASD-like traits by affecting the central release of AVP and/or OXT, which can impact social behavior, from either somatodendritic compartments (Chini et al., 2017) and/or axonal projections of PVN OXT-ergic neurons to extrahypothalamic social centers (Zhang et al., 2021). Moreover, such PBDE effects on the HNS and social behavior may, in part, be mediated through PBDE-induced TH disruption. Indeed, much evidence exists on endocrine disruption of thyroid (and reproductive) endocrine systems by PBDEs (Costa & Giordano, 2007; Zhao et al., 2015). However, further research is needed to shed light on these interrelated processes as potential mechanisms of PBDE action.

To the best of our knowledge, no comprehensive characterizations of ASD neurobehavioral phenotypes in murine models of developmental BDE mixtures or single congener exposures exist. We, therefore, conducted a study which comprehensively tested the effects of maternal DE-71 on the core ASD domains and constructs that represent co-morbidities in ASD, namely cognitive impairment, hyperactivity, anxiety, depression and sensorimotor integration. The present study builds on our prior work in female offspring and extends the finding to male mice (Kozlova et al., 2021). Since ASD affects each sex differently and PBDE congeners show estrogenic, anti-estrogenic and/or androgenic actions (Ren & Guo, 2013), it is important to characterize the effects of PBDEs in each sex. In this study we exposed mouse dams to the commercial PBDE mixture, DE-71, at low doses designed to model chronic, low-level exposure to translationally relevant congeners. We show that perinatal exposure to DE-71 leads to enduring impairments in social recognition and general memory, repetitive behavior, olfactory function, and prosocial neuromolecular markers in brain regions involved in regulating complex social behaviors. Our findings emphasize the importance of considering sex as a biological variable in studies of environmental neurotoxicity and implicate PBDEs as contributors to ASD-related neurodevelopmental outcomes.

## Methods

### Animal Housing and Care

C57Bl/6N mice were generated using breeders originally obtained from Charles River Labs (West Sacramento, CA). Mice were housed 2-4 per cage in standard static polycarbonate plastic cages with corn-cob bedding in a specific pathogen free vivarium and kept on a 12:12-h light:dark cycle in a controlled temperature (21.1–22.8°C) environment. Relative humidity ranged between 20% and 70%. Mice were provided rodent chow (LabDiet 5001; Laboratory Diets, USA) and water *ad libitum*. Care and treatment of animals was performed in compliance with guidelines from and approved by the University of California Riverside Institutional Animal Care and Use Committee (AUP#5, 20170026 and 20200018).

### Experimental Design and DE-71 Exposure

DE-71 (technical penta-bromodiphenyl oxide; Lot no. 1550OI18A), was obtained from Great Lakes Chemical Corporation (West Lafayette, IN). DE-71 dosing solutions were prepared in corn oil vehicle (VEH/CON) at two concentrations: 0.1 mg/kg/day (L-DE-71) and 0.4 mg/kg/d (H-DE-71) using 2 mL of stock solution per kg body weight. These doses were selected to match the molar concentrations of BDE-47 used in previous mouse studies (Woods et al., 2012); (Wang et al., 2018).

Offspring were exposed to DE-71 via maternal transfer through a 10-week dosing regimen (**Fig.1a**) as described previously (Kozlova et al., 2020). Mice were randomly assigned to one of the three exposure groups: corn oil vehicle control (VEH/CON), L-DE-71 or H-DE-71. After 3 weeks of pre-dosing, virgin females were paired with an untreated male using harem-style breeding. F1 offspring were weaned at postnatal day (PND) 21 and housed in same-sex cages (2-4/cage). Adult male offspring (F1) were subjected to behavioral testing and later sacrificed by exsanguination via cardiac puncture under terminal isoflurane anesthesia (5%) followed by cervical dislocation.

To reduce cross-over effects, behavioral tests were distributed across three different cohorts. Mice were run through a battery of behavioral tests in the following order for Cohort 1 (mean age): Suok (PND 43); social novelty preference test (SNP; PND 71); 3 chamber sociability (PND 87); elevated plus maze (EPM; PND 72) and open field test (OFT; PND 100). The brains of Cohort 1 were collected at sacrifice on PND 110 and analyzed via RT-qPCR. The following tests were performed on Cohort 2: Marble burying (MB; PND 81); olfactory habituation/dishabituation (OHT; PND 79); olfactory preference (OPT; PND 102); forced swim test (FST; PND 74). Cohort 3 was subjected to social memory recognition (SRM; PND 30); juvenile OFT (PND 31); juvenile MB (PND 35); novel object recognition (NOR; PND 111) tests. Analytical characterization by mass spectrometry was performed on brains from Cohort 1 (PND 110) and a subset of Cohort 3 (PND 30). Enzyme-linked immunosorbent assays (ELISA) were performed on plasma from Cohorts 1-3. Whenever possible, the dam or litter was used as the statistical unit of analysis for F1 (i.e., values of offspring in each litter were averaged to represent one biological replicate). In addition, results were replicated in a minimum of 3 independent experiments.

### PBDE Congener Analysis in Perinatal Offspring Brain via High Resolution Gas Chromatography–High Resolution Mass Spectrometry (HRGC/HRMS)

Using High Resolution Gas Chromatography–High Resolution Mass Spectrometry (HRGC/HRMS), PBDE concentrations were measured in PND 30 whole brain homogenates (0.1-0.2 g) as described (Z.-M. Li et al., 2020) by the Schramm laboratory at Helmholtz Zentrum München. PBDE analytes included 37 PBDE congeners (BDE-7, 10, 15, 17, 28, 30, 47, 49, 66, 71, 77, 85, 99, 100, 119, 126, 138, 139, 140, 153, 154, 156, 176, 180, 183, 184, 191, 196, 197, 201, 203, 204, 205, 206, 207, 208, 209). Samples were ground and homogenized to a fine powder under liquid nitrogen. Each sample (100-200 mg) was mixed with CHEM TUBE-Hydromatrix (Agilent Technologies) and spiked with ^13^C-labelled PBDE standard mix (BFR-LCS, Wellington Laboratories). For pressurized liquid extraction (Dionex ASE 200), n-hexane/acetone (3:1, v/v) was used at 120°C and 12 MPa. The volume of the extract was reduced to ∼5 mL using a vacuum rotary evaporator. Samples were purified using an automated system (DEXTech, LCTech, Germany), where the sample was passed and fractionated over an acidic silica, alumina and carbon column. Concentrated extracts were spiked with the recovery standard (BFR-SCS, Wellington Laboratories) and analyzed by HRGC/HRMS (Agilent 6890/Thermo MAT95 XL) using electron impact ionization (EI), in selected ion monitoring mode. The instrumental parameters are listed in **Supplementary Table 1**. Average recovery for ^13^C-labelled PBDE standards ranged between 40 and 120%. All samples were blank-corrected on a congener-specific basis using the average of three procedural blank samples. Analytes with concentrations after blank correction that were lower than three times the standard deviation of the blank values or that were not detected before blank correction were considered as not detectable (n.d.). The limit of quantification (LOQ) of the instrumental methodology was considered as a signal/noise ratio of 9:1 (**Supplementary Table 2**). Congener concentrations that were below the detection limit were assigned a randomly generated value of LOQ/2. The accuracy of our method was confirmed by successful participation in interlaboratory comparison studies.

### PBDE Congener Analysis in Adult Offspring Brain via Gas Chromatography in tandem with electron capture negative ion mass Spectrometry (GC/ECNI/MS)

PBDE concentrations were measured in PND110 whole brain homogenate extracts by the Stapleton laboratory at Duke University using gas chromatography coupled with electron capture negative ion mass spectrometry (GC/ECNI-MS; Agilent 5975N MS) as described previously (Kozlova et al., 2020). Briefly, approximately 0.2-0.5 grams of tissue were first ground with clean sodium sulfate, spiked with two isotopically labeled standards (F-BDE-69 and 13C BDE-209) and then extracted using 50:50 DCM:hexane. Extracts were concentrated, measured for lipid content using a gravimetric analysis, and then purified using acidified silica prior to analysis for 26 different PBDE congeners ranging from BDE-30 to BDE 209. Laboratory processing blanks (clean sodium sulfate only) were analyzed alongside samples to monitor background contamination. Recoveries of F-BDE-69, and 13C BDE 209, averaged 91 (+/− 6.9%) and 106 (+/− 19.9%), respectively, in all samples. All samples were blank-corrected on a congener-specific basis using the average concentrations measured in the laboratory processing blanks. Method detection limits (MDLs) were estimated using either a signal to noise ratio of 10, or, if analytes were detected in laboratory blanks, by calculating three times the standard deviation of the laboratory blanks. MDLs differed by congener and ranged from 0.8 (BDE-47) to 6.6 ng/g (BDE-206).

### Neurobehavioral Testing Paradigms

At least 30 min prior to testing, mice were moved to a designated behavior room. Ethanol (70%) was used to remove debris and odors between individual mouse trials. Unless stated otherwise, mouse behavior was scored using automated video-tracking software (Ethovision XT 15, Noldus) or manual scoring software (BORIS (Friard & Gamba, 2016) or JWatcher), performed blind to treatment by trained observers. Mice were tested between 10 am and 4 pm during the light phase under bright light conditions, unless otherwise stated.

### Social Novelty Preference

Social novelty preference (SNP) was conducted and analyzed according to methods adopted from published protocols (Moy et al., 2004). Briefly, mice were habituated for 30 min to a polycarbonate cage identical to their home cage (26 cm length x 16 width cm x 12 cm height), followed by 30 min to two wire interaction corrals (11 cm height × 10 cm diameter) placed on each side of the cage. During a 5-min training trial, a stimulus mouse was placed into one corral while the empty corral was removed. After a 30 min retention period, social recognition was assessed in the following 5 min test, during which the test mouse explored the same stimulus mouse (now familiar mouse) versus a novel stimulus mouse. Prior to testing days, sex- and age-matched conspecific stimulus mice were trained to stay in corrals for 15 min for 3 times per day for 7-14 d. Stimulus mice were single-housed in order to preserve their unique scent. Investigation by test mouse was measured as time spent sniffing (snout within 2 cm of stimulus). Test robustness was measured using an Investigation Index calculated as the ratio of time spent investigating the novel mouse to total investigation time during the training trial (**Supplementary Figure 2**). Social recognition is represented as decreased investigative behavior toward a conspecific rodent during a second encounter and relatively more time spent investigating novel stimulus as percent of total investigation time. To evaluate between group differences, a Discrimination Index (DI) was calculated as the ratio of time (T) spent investigating Novel - Familiar/total investigation time in test period (DI = (T_Novel – T_Familiar) / (T_Novel + T_Familiar)).

### Social Recognition Memory Test

A two-trial social recognition memory test (SMRT) was performed as previously described (Kogan et al., 2000), (Tanimizu et al., 2017) to assess long-term social recognition memory. Test mice (PND 28-40) were exposed to a juvenile sex-matched conspecific stimulus mouse (PND 15-32) during two 3 min trials following an intertrial delay of 24 h. For each experiment, test mice were individually placed into polycarbonate cages (27 x 16 x 12 cm) and allowed to habituate for 1 h under dim conditions. A juvenile sex-matched conspecific was then placed into the cage and the mice were allowed to interact for 3 min (Trial 1). In Trial 2, performed 24 h later, the same test mouse was exposed to either the familiar (stimulus from Trial 1) or a novel stimulus. Each stimulus was not used more than 4 times. The tests were digitally recorded and scored for social investigation behavior by a blind trained observer including direct contact (including grooming, licking, and pawing) with the juvenile while inspecting any part of the body surface, sniffing of the mouth, ears, tail, ano-genital area, and close following (within 1 cm) of the juvenile (Thor, 1982). Only animals that reliably investigated the juveniles without displaying any aggressive behavior were retained for Trial 2 (∼85% of the original population). To evaluate the differences in ability to form a long-term social memory a Recognition Index was calculated as the ratio of the duration of investigations on Day 2 and Day 1 (RI = T_Day2 / T_Day1).

### Three-Chamber Sociability

A sociability test was performed in a three-chamber apparatus using methods described previously by (Yang et al., 2011). The day of the experiment the test mouse was allowed to acclimate to the apparatus for 20 min before testing – 10 min in the central chamber with the doors closed, followed by 10 min in the entire empty arena with the doors open. The subject was then briefly confined to the center chamber while empty inverted wire corrals (non-social stimulus) were placed in the side chambers. A male stimulus mouse, used as the social stimulus, was placed inside one of the corrals. After both stimuli were positioned, the testing began when doors were reopened and the mouse was allowed to access all three chambers for 10 mins. Number of entries and time spent in the chambers as well as sniffing time were manually determined from recorded videos using the event-recorder software JWatcher. Sniffing time, measured as time test mouse snout was within 2 cm of enclosed mouse, has been suggested to have superior validity over chamber time scores since active behaviors that are most directly related to social investigation are captured (Fairless et al., 2011) and the physical proximity allows for transmission of volatile and nonvolatile odorants (Brennan & Kendrick, 2006); (Luo et al., 2003); (Fairless et al., 2011).

### Marble Burying and Nestlet Shredding

The Marble Burying (MB) and Nestlet Tests were utilized for analysis of elicited repetitive behavior in rodents that is analogous to those observed in individuals with autism (Silverman et al., 2010). During the marble burying test, the test mouse was placed in the corner of a polycarbonate cage (19 x 29 x 13 cm) containing bedding at 5-cm deep as described (Angoa-Pérez et al., 2013). The mouse was then allowed to interact for 30 min with an equidistantly aligned marble array (8 x 4 used for adults or 5 x 4 used for juveniles). A photograph of the cage was taken prior to and after the test and images were scored by 2-3 investigators who were blind to treatment. A minimum ⅔ of the marble was defined as being buried in the 32-marble array and ½ buried in the 20 array. The number of buried marbles was then averaged between the scorers (**Supplementary Figure 3**). After a 5 min rest period, the Nestlet shredding test was performed. The test mouse was placed into another cage of the same size with 0.5 cm of bedding containing a pre-weighted square of cotton fiber Nestlet. After 30 min, the remaining Nestlet mass was weighed and percent Nestlet shredded was calculated.

### Novel Object Recognition Test

The novel object recognition test (NORT) was used to assess *non-social* recognition memory. A two-day protocol with a short- and long-term retention time was used. On Day 1, the test mouse was habituated to an empty square Plexiglas open field arena (39 x 39 x 38 cm) for 15 min as described (Murai et al., 2007), followed by a 20 min rest in its home cage. During the acquisition phase, the test mouse was placed in the open field containing two identical objects (F vs F’) and allowed to freely explore the environment and objects. During the short-term memory (30 min retention) testing session, the test mouse was again placed in the apparatus and allowed to explore a familiar and novel object (F vs N). After a 24 h retention time (Day 2), long term memory was assessed by placing mice into the open field containing both the familiar and a new novel object (F vs N’). All test/train sessions lasted 5 min. Preference for the novel object was expressed as the ratio of time exploring the novel stimulus relative to the total exploration time. To evaluate the differences in ability to form NOR memory, a Discrimination Index was calculated as the difference in exploration time between novel and familiar objects relative to total exploration time, where 0 indicates equal preference (DI = (T_Novel – T_Familiar) / (T_Novel + T_Familiar)). Test object pairs were first validated for lack of intrinsic preference in untreated mice. Data was analyzed using Ethovision.

### Innate Olfactory Preference Test

To test the ability to detect attractive or aversive odorants, the innate Olfactory Preference Test (OPT) was performed and analyzed as described (Kobayakawa et al., 2007), (Witt et al., 2009). Mice were acclimated to a large empty test cage (19 x 29 x 13 cm). After 15 min mice were sequentially transferred to three other cages every 15 min each containing a filter paper (2 x 2 cm) treated with 500 uL of a test odorant: 10% peanut butter, 1% vanilla, 1% butyric acid, or deionized water. The four test odorants were presented to the test mouse in a randomized order. Time spent sniffing the filter paper during the 3-min odorant trials was video-recorded and later measured.

### Olfactory Habituation Test

The Olfactory Habituation/Dishabituation test (OHT) was used to assess mice’s ability to detect and differentiate social and non-social odorants (Arbuckle et al., 2015), (Silverman et al., 2010). Mice were acclimated for 45 min to an empty cage with a cotton-tipped applicator inserted through the water bottle hole. Non-social odors were prepared from extracts immediately before testing. They included: (1) deionized water; (2) almond (1:100 dilution); (3) banana (1:100) (McCormick). Two social odors were obtained the morning of test day by swiping applicator across the bottom of two different stimulus cages containing soiled bedding from sex-matched conspecifics which had no previous contact with the test subject. These cages house 3-4 mice and bedding was at least 3 d old. Stimuli were presented in the following order: water x 3, almond x 3, banana x 3, social odor 1 x 3, social odor 2 x 3. Time spent sniffing the presentation of the test applicator during the 2-min trial periods was recorded on a stopwatch. Habituation, defined as a decrement in olfactory investigation of the same odor after repeated presentations and dishabituation, defined as a reinstatement of olfactory investigation upon presentation of a new odorant were assessed.

### Elevated Plus Maze

Anxiety was assessed using the elevated plus maze (20 x 22 cm; elevated 92 cm) (EPM), constructed from black Plexiglass as described (Lister, 1987). Two open arms and two closed arms received differential lighting, i.e., 300 and 30 LUX, respectively, to create anxiogenic (open) and anxiolytic conditions (closed). At the beginning of each trial, the test mouse was placed in the middle of the apparatus facing the open arm and was allowed to explore the apparatus for a total of 5 min. Activity was monitored by an overhead digital camera and video recordings were analyzed for time spent in each arm and the frequency of entries using BORIS.

### Forced Swim Test

The forced swim test (FST) was used to assess depressive-like behavior using an apparatus consisting of a vertical plexiglass cylinder (127 X 305 mm) filled to a depth of 150 mm with water maintained at 23-25 (Lucki et al., 2001). Testing was conducted under dim room lighting conditions with an overhead light placed above the cylinders. Mice were placed individually into the water column for a 6 min test trial. After the test, mice were removed and dried before returning to their home cage. Swimming behavior was analyzed during the last 4 min using Ethovision. Active swimming was considered as time spent struggling/paddling with more than one limb. Drifting or passive movements used to maintain verticality in the water column were not considered swimming and instead were used to record time spent immobile.

### Suok

Suok is an elevated platform behavioral paradigm used to analyze anxiety, anxiety-induced motor impairments and motor-vestibular anomalies in mice (Kalueff and Touhimaa, 2005a). The apparatus consists of a smooth aluminum rod (2 m long, 3 cm diameter) elevated to 20 cm and fixed to two clear acrylic walls as described (Kalueff et al., 2008). Bilateral to a central segment (38 cm) of the aluminum rod are 10 cm segments labeled by line markings visible to the experimenter. After acclimation to the dimly lit testing room for 30-60 min, several behaviors are scored over a 5 min trial: (1) horizontal and locomotor (normalized) activity - assessed by number of segments traveled, (2) sensorimotor coordination -measured by the number of hind leg slips and falls from the rod, (3) exploratory behavior like side looks and head dips, (4) anxiogenic behaviors such as increased latency to leave the central zone and unprotected stretch-attend postures, in which the mouse stretches forward and retracts without moving its feet, considered a non-social form of ambivalence (Kaesermann, 1986); (Benneh et al., 2018), (5) vegetative responses (combined number of urinations and defecation boli), and (6) autogrooming behaviors. Hyperactivity, loss of sensorimotor coordination, increased anxiety and displacement behavior are represented by elevated values for #1, 2, 4 and #5, and 6 (Kalueff & Tuohimaa, 2005), (Benneh et al., 2018). Measures were recorded manually by stopwatch. Locomotor activity was calculated as total test time minus time spent immobile in the center.

### Open Field Test

The open field test allows rapid assessment of rodent locomotion, anxiety and habituation without a training requirement (Hall, 1934). The open field apparatus, a plexiglas square arena of 39 x 39 x 37.8 cm, was designed as a large, brightly lit, open and aversive environment for test rodents. Mouse activity over a 1 h period was digitally recorded and scored using Ethovision for distance traveled, velocity and total time in periphery (10 cm adjacent to wall) and center. Time in the periphery represents non-anxious motor activity since it is positively correlated with frequency of entries into open and closed arms on the elevated plus maze, and is negatively correlated to defecation (Carola et al., 2002). The latter represents anxious motor activity (Seibenhener & Wooten, 2015). Reduced distance traveled over the 1 h period also indicates habituation to novelty and learning (Bolivar et al., 2000).

### RNA Extraction From Brain Micropunches

Animals were sacrificed by exsanguination via cardiac puncture shortly after PND 110. Whole brains were rapidly dissected and flash frozen in 2-methylbutane (Fisher chemical) over dry ice and stored at −80 ^0^C until further use. Cryosections (0.3-0.5 mm thick) of mouse brain were generated and mounted on sterile glass slides and frozen followed by storage at −80^0^C. Five regions of interest were punched out bilaterally from tissue sections under a stereomicroscope using a handmade microdissecting needle (16-gauge) adapted from the Palkovits micropunch technique (Palkovits, 1973). The anatomical precision of samples was based on the atlas of Paxinos and Franklin. Tissue punches were immediately homogenized in TRIzol Reagent (Thermo Fisher Scientific, USA) using a hand-held homogenizer, snap frozen over dry ice and stored at −80^0^C until further use. Total RNA was isolated via a modified partial phenol-methanol extraction protocol using the RNeasy Micro Kit (Qiagen, USA). Purity and quality of RNA were assessed photometrically using the ratio of optical density (OD) at 280 nm over 260 nm (NanoDrop ND-2000). RNA integrity was assessed using an Agilent 2100 Bioanalyzer.

### Quantitative polymerase chain reaction

All molecular work was carried out in adherence to MIQE guidelines (Bustin et al., 2009) and previously described (Kozlova et al., 2021). Oligonucleotide PCR primers were custom designed and synthesized, or ordered as predesigned assays from Integrated DNA Technologies. Primers were designed to meet several criteria using NCBI Primer Blast and then optimized by testing against complementary DNA generated using RT-PCR and gel electrophoresis. Only primers that gave single-band amplicons in the presence of reverse transcriptase (RT) and that matched the base length of the predicted target were chosen for further optimization via qPCR. Selected primers were then optimized to yield 90% to 110% efficiency. *Oxtr* and its reference gene, *ActB*, were multiplexed using hydrolysis probes with double-quenchers. For all other primers, intercalating dye chemistry was used. Primer sequences have been previously reported (Kozlova et al., 2021). RT-qPCR was performed on a CFX Connect thermocycler (Biorad, Hercules, CA) with the Luna Universal or Probe one-step qPCR Master Mixes (New England Biolabs, Ipswich, MA). RNA (1-4 ng) was used per reaction run in triplicate. In each experiment, no-template controls (NTCs) without mRNA were run to rule out extraneous nucleic acid contamination and primer dimer formation. Negative RT controls, which contained the complete RNA synthesis reaction components without the addition of the RT were used to rule out presence of genomic DNA (gDNA). Fold-change gene expression was measured relative to the reference gene, *ActB*, and differential gene expression was determined compared to null group (VEH/CON) using the Pfaffl method (Pfaffl, 2001).

### Enzyme Immunoassays

Blood was collected by cardiac puncture and the plasma separated at 2000 x g centrifugation for 20 min at 4. Plasma levels of the neuropeptides OXT and arginine8-vasopressin (Arg^8^) were quantified using commercially available ELISA kits from Arbor Assays (Ann Arbor, MI USA Ref: OXT, K048-H1, Arg^8^, K049-C1), Enzo Life Sciences (Farmingdale, NY USA, OXY: ADI901153A0001, Arg^8^: ADI-900-017) and Invitrogen (Waltham, Massachusetts, OXT, EEL139) following the manufacturer’s instructions. For the Arbor Assay kits, in order to reduce the non-specific binding, samples were first treated using the acetone-based extraction solution followed by vacuum lyophilization of the resulting supernatant. For OXT, the colorimetric reaction product was read as optical density at 450 nm on a plate reader (SpectraMax 190, Molecular Devices). The kit has a sensitivity of 1.7 pg/mL in a dynamic range of 16.38-10,000 pg/mL. ARG^8^-Vasopressin was detected using a luminescence plate reader (Victor3, Perkin Elmer). The ARG^8^-Vasopressin kit has a sensitivity of 0.9 pg/mL in a dynamic range of 1.638-1,000 pg/mL. For the Enzo Life Sciences kits, samples underwent solid phase extraction using 200 mg C18 Sep-pak columns as previously described (Deol et al., 2020). The invitrogen ELISA has a sensitivity of 9.38 pg/mL in a dynamic range of 15.63-1,000 pg/mL. Plasma oxytocin and arginine vasopressin were quantified by interpolating absorbance or luminosity values, respectively, using a 4-parameter-logarithmic standard curve (MyAssays).

### Statistical Analyses

Statistical analysis was performed in GraphPad Prism (version 8.4.3 San Diego, CA, USA). The normality of data distribution was evaluated with the Shapiro-Wilk test and the equality of variances across groups was compared using the F-test. For parametric data, one-tailed unpaired or two-tailed paired Student’s *t*-tests were used to analyze congener levels as well as performance in the SNP, SMRT, and NOR tests. One-way ANOVA followed by Tukey’s or Dunnett’s *post hoc* tests was used to compare means across groups for all other data involving three-group comparisons. A Brown-Forsythe ANOVA or Welch’s correction were used instead if the group variances were significantly different. For nonparametric data, a Kruskal-Wallis H test with Dunn’s post hoc was used to compare differences across groups. A one-sample t-tests were used to determine whether the indexes on social tests and NORT differed significantly from the hypothetical values as described. Type 1 error rate (α) was set at 0.05; F and *P* values are presented in the figure legends or in Supplementary files. The data are expressed as the mean ± s.e.m, as mean with individual values as ‘before-after’ bars or as median and inter quartile range representing minimum and maximum values in whisker plots. Statistical outliers were excluded when values exceeded 2 x standard deviation from the mean. Technical outliers were excluded when animals were unable to perform behavioral tests. To examine interrater reliability on marble burying and nest scores, the Bland-Altman method was used to calculate bias as the mean of the differences (0 means two judges are not producing different results) and precision as 95% limits of agreement (standard deviation of mean bias +/− 1.96).

## Results

### DE-71 Dosing Paradigm and Maternal Parameters

C57Bl/6 dams were exposed to DE-71 and their F1 male offspring were investigated (**Fig. 1**). Using this dosing paradigm, we have previously reported neurodevelopmental delays that could not be attributed to differences in litter size at birth, secondary sex ratio, gestational maternal parameters nor maternal retrieval behavior (Kozlova et al., 2020), (Kozlova et al., 2025). Therefore, our current results are interpreted without cause assigned to maternal behavior deficits. In this study, offspring body weights measured at PND 44 were not affected by treatment(**Supplementary Figure 1**). This is consistent with no differences in male body weight or composition in adulthood that we previously reported (Kozlova, Denys, et al., 2022).

**Fig. 1.**
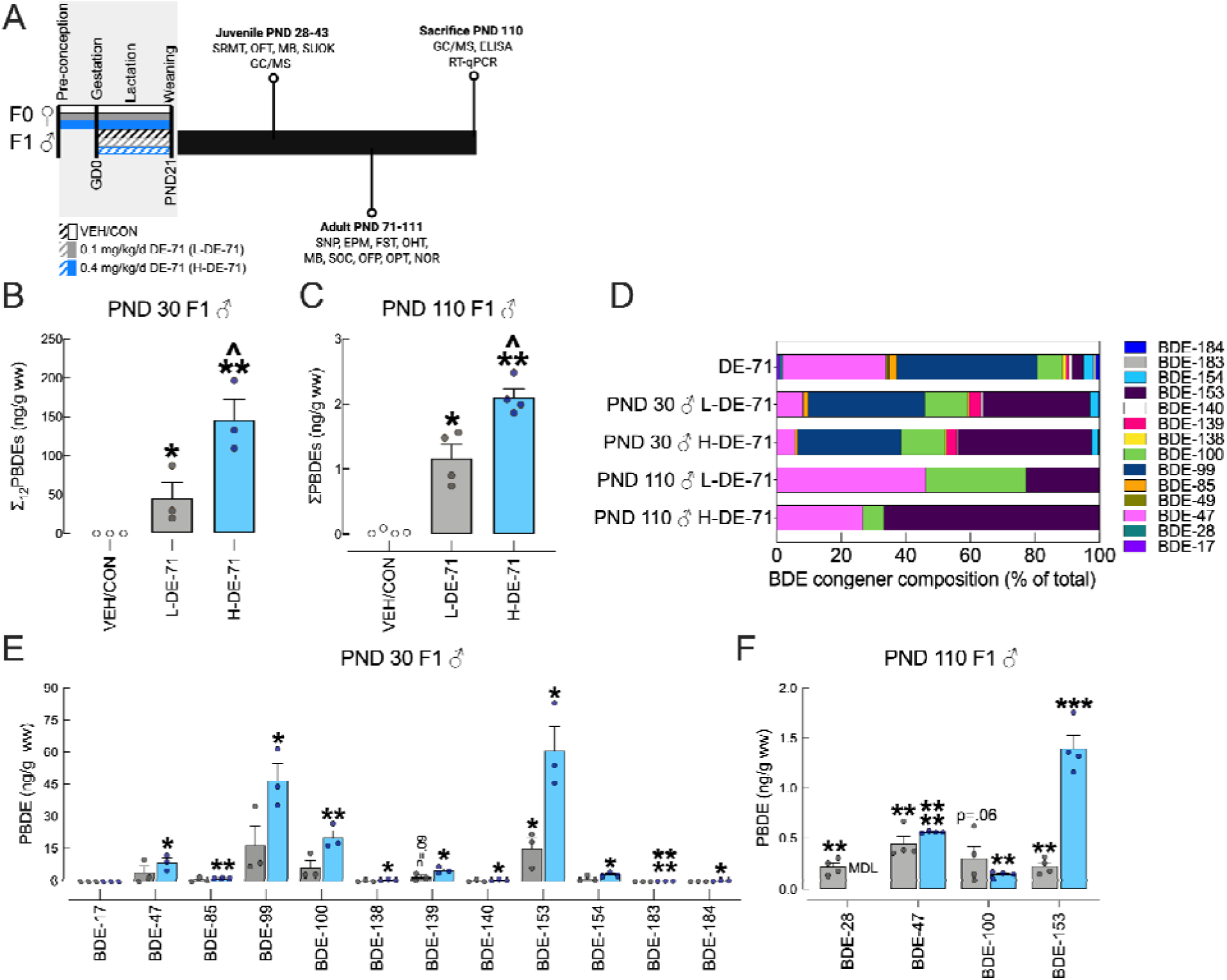
Maternal dosing paradigm for DE-71 produces persistent BDE congener penetration in male F1 offspring brain. **a** Direct exposure to DE-71 in adult dams (F0♂; solid shading), began ∼3-4 weeks pre-conception and continued during lactation until pup weaning at PND 21. Indirect exposure in male offspring (F1♂; hatched shading) occurred perinatally (GD 0 to PND 21). Offspring were subjected to behavior tests at different ages as shown. **b** The sum concentrations of the 12 PBDE congeners (∑_12_PBDEs) in ng/g wet weight (ww) detected at PND 30. **c** The sum concentrations of the 3-4 PBDE congeners (∑PBDEs) detected in ng/g ww at PND 110. **d** BDE composition (% total) in DE-71 and in offspring brain samples. The 7 congeners that comprise <1% of DE-71 were displayed as 1% for better distinction. **e-f** Absolute congener concentrations. *significant difference compared to VEH/CON, \**P*<.05, \*\**P*<.01, \*\*\**P*<.001, \*\*\*\**P*<.0001; ^significant difference compared to L-DE-71, ^*P*<.05. *n*, 3-4 individuals/group. EPM, Elevated Plus Maze; ELISA, Enzyme-Linked Immunosorbent Assay; FST, Forced Swim Test; GD, gestational day;vMB, Marble Burying; NOR, Novel Object Recognition; OFT, Open Field Test; OHT, Olfactory Habituation/Dishabituation Test; OPT, Olfactory Preference Test; PND, postnatal day; RT-qPCR, Reverse Transcription Quantitative Polymerase Chain Reaction; SNP, Social Novelty Preference Test; SRMT, Social Recognition Memory Test

### PBDE Congener Analysis in Offspring Brain

PBDE congener content was determined using HRGC/HRMS or GC/ECNI-MS in brains from F1 exposed male juveniles soon after weaning (PND 30) or as adults (PND 110), respectively. Thirty-seven BDE congeners representing di-through deca-BDEs were analyzed since commercial DE-71 mixtures contain primarily tetra-, penta-, and hexa-brominated congeners and traces of tri- and hepta-brominated congeners (LaA Guardia et al., 2006). Raw values by lw and/or ww are listed by exposure group in **Supplementary Tables 3, 4, 5.** In **Figure 1b** mean group values of sum concentrations of 12 PBDEs detected (∑_12_PBDEs) indicates penetration of L-DE-71 (*P<*.05) and H-DE-71 (*P*<.01) into PND 30 offspring, relative to VEH/CON, confirming that the dosing regimen led to maternal transfer of PBDEs to offspring brain in a dose-dependent manner (*P*<.05). Mean ∑_12_BDE values were 45.4 and 147 ng/g ww for L-DE-71 and H-DE-71, respectively (**Fig. 1b**).

Collectively, 6 of 12 congeners (BDE-47, −99, −100, −139, −153, −154) in L-DE-71 and H-DE-71 accounted for 97% of all PBDEs penetrating the brain at PND 30. These same 6 congeners comprise 97% of the DE-71 mixture (**Fig. 1d**). The remaining congeners in our samples (BDE-17, −85, −138, −140, −183 and −184) made up 2.8 and 2.0%. However, when compared to VEH/CON, mean concentrations for only BDE-153 (*P*<.05) and −139 (*P*=0.09) in L-DE-71 were comparatively different perhaps due to the low dose and sample size. In contrast, the mean concentrations for all 6 major BDE congeners were significantly greater in H-DE-71 vs VEH/CON (*P*<.05-.0001). Of the 6 minor congeners, 5 (i.e. BDE-85, −138, −140, −183, −184) were significantly greater in H-DE-71 vs VEH/CON (*P*<.05-.0001). Notably, BDE-153 was 12-fold enriched and BDE-47 was ∼6-fold reduced relative to the DE-71 mixture as reported previously (Kodavanti et al., 2010) (**Fig. 1e**). Both BDE-153 and 47 are reported to be transferred (Vizcaino et al., 2011).

While the overall magnitude decreased ∼75-fold from PND 30 to PND 110, i.e., 1.2 and 2.1 ng/g ww (78.6 and 230 ng/g lw) for L- and H-DE-71, respectively, the mean group values for ∑PBDEs show similar penetration trends of L-DE-71 (*P<*.05) and H-DE-71 (*P*<.01) at PND 110 (**Fig. 1c**). At PND 110 only 3-4 BDE congeners were detected in L-DE-71 (BDE-28/33, −47, -100, −153) and H-DE-71 (BDE-47, −100, −153) (**Fig. 1f**). The mean concentrations for 3 of the major BDE congeners in L-DE-71 were significantly greater (BDE-28, −47 and −153, *P*<.01) or apparently greater than in VEH/CON (BDE-100, *P*=.06). For H-DE-71 mean concentrations were greater for BDE-47, −100, −153 (P<.01-.0001). **Figure 1d** depicts BDE congeners by percent composition.

### Perinatal exposure to DE-71 Induced ASD-like deficits in F1 male

#### Social Novelty Preference

Testing mice on a social novelty preference (SNP) test has been suggested to be ethologically relevant to symptoms observed in autistic individuals (Moy et al., 2004). On this test, the VEH/CON offspring but not the L-DE-71- or H-DE-71-exposed mice showed a preference for the novel over familiar stimulus (**Fig. 2a**, *P*<.01). Within group comparisons of mean recognition index (RI) scores showed a significant increase from “0” (a hypothetical value indicating no preference) in VEH/CON but not DE-71 exposed males (**Fig. 2b**, *P*<.01, one sample t-test). A one-way ANOVA revealed differences as compared to VEH/CON for L-DE-71 (apparent, *P*=.08) and H-DE-71 (*P*<.05). Investigation index scores comparing investigation on Trial 2 vs Trial 1, a measure of sociability, indicated that the reduced exploration of a novel versus familiar stimulus on SNP by L-DE-71- and H-DE-71-exposed offspring was not due to a lack of social interest (**Supplementary Figure 2**).

**Fig. 2.**
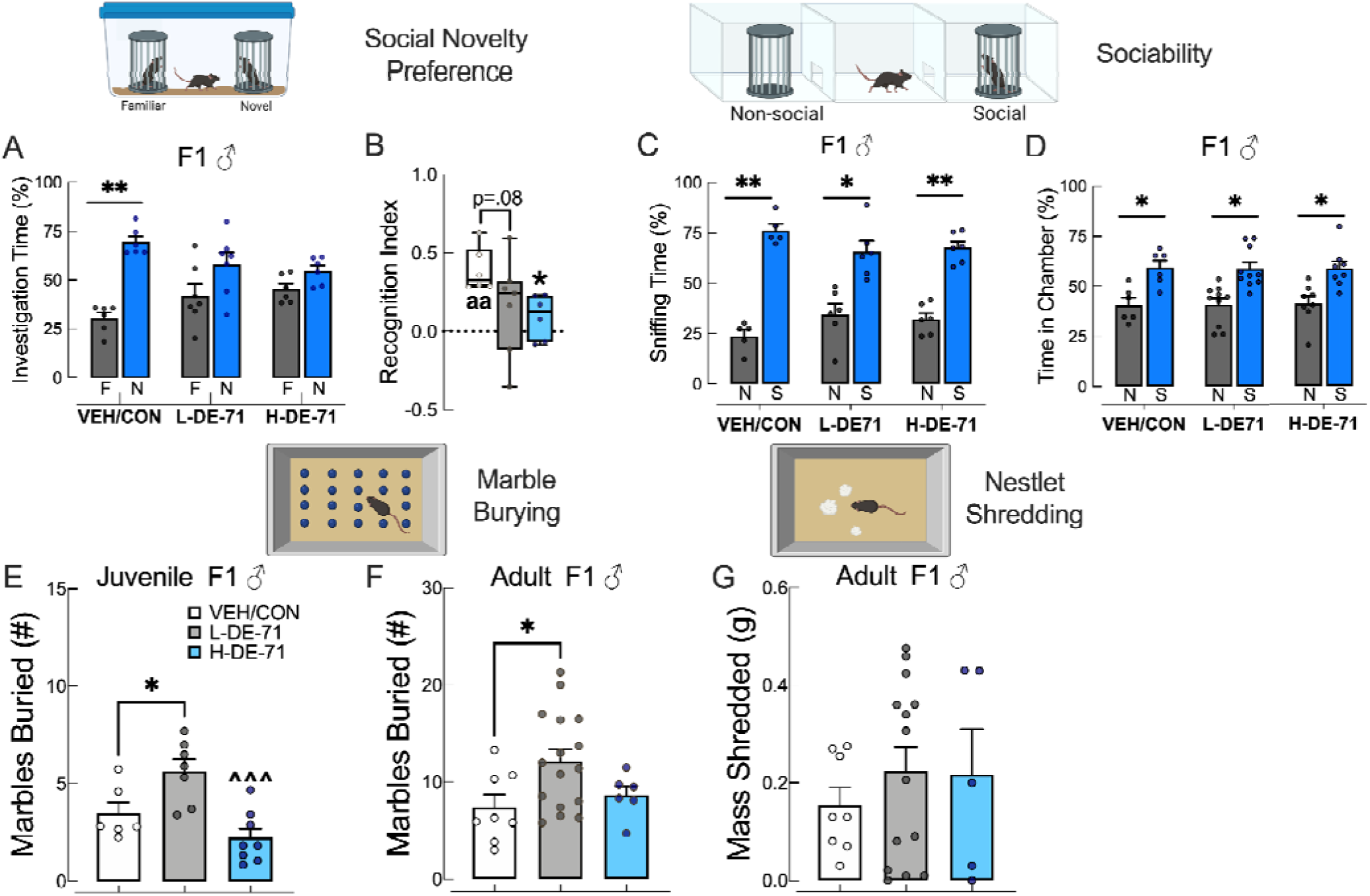
Early-life exposure to DE-71 induces deficits relevant to core symptoms of ASD in juvenile and adult F1 males. **a** Social Novelty Preference scores for male offspring: unlike the VEH/CON, L-DE-71 and H-DE-71-exposed male offspring failed to prefer a novel (N) relative to a familiar (F) conspecific stimulus. **b** Recognition Index scores showed that only VEH/CON were significantly different from “0” (a hypothetical score representing no preference), indicating proper social recognition. **c** Time spent sniffing with a social (S) vs non-social (N) stimulus on the Sociability test. All exposure groups spent significantly more time sniffing social stimulus, indicating normal sociability. **d** Chamber time scores in Sociability test. All groups showed significantly greater time spent with S relative to N. **e** Marble Burying scores in juvenile offspring showed that DE-71 exaggerated digging behavior in a dose-dependent manner. **f** Marble Burying scores in adult offspring showed similar results. **g** Nestlet shredding was not affected in exposed offspring. *significantly different from familiar (a) or from non-social stimulus (c,d) or from VEH/CON (b,e,f,g), \**P*<.05, \*\**P*<.01, ^a^significantly different from “0”, ^aa^*P*<.01, one-sample t-test. *n*, 6-7 litters/group (a-b), 5-10 litters/group (c, d), 6-8 litters/group (e), 6-16 litters/group (f), 5-14 litters/group (g). F, familiar; N, novel (a), non-social (c,d); S, social

#### Sociability

To determine social interest, an independent social cognition domain, we examined mouse behavior on a 3-chamber sociability test. All F1 groups showed preference for a novel social stimulus relative to a non-social novel stimulus as measured by sniffing time (**Fig. 2c**, *P*<.05, <.01), as well as time in chamber, both indicating normal sociability (**Fig. 2d**, *P*<.05). As a measure of test robustness, there was no indication of side preference during training (**Supplementary Figure 2**).

#### Repetitive Behavior

On the marble burying test, which measures repetitive and perseverative behavior in rodents (Thomas et al., 2009), L-DE-71 (but not H-DE-71) juvenile PND 30 male offspring buried a significantly greater number of marbles relative to VEH/CON (**Fig. 2e,** *P*<.05). There was also a significant dose effect of DE-71 since H-DE-71 males buried fewer marbles than L-DE-71 males (**Fig. 2e,** *P*<.001). Similar results were found for a subgroup of exposed male offspring tested in adulthood (**Fig. 2f,** *P*<.05). There were no group differences found on a nestlet shredding test (**Fig. 2g**).

### Perinatal exposure to L-DE-71 but not H-DE-71 reduced long-term social recognition memory

We determined that SNP scores requiring a 30 min memory retention were abnormal in exposed F1 males. To measure group effects on *long-term* social recognition memory, we subjected mice to a social recognition memory test (SRMT) (Kogan et al., 2000; Tanimizu et al., 2017). On this test, mice with intact memory exhibit less time investigating a familiar juvenile conspecific 24 h after a first encounter of 3 min of Day 1 (Kozlova et al., 2021). **Figure 3a** shows that VEH/CON and H-DE-71 mice were able to form a memory for a previously encountered mouse, since they spent significantly less time with the familiar stimulus mouse on Day 2 vs Day 1 (*P*<.01 and *P*<.05). In contrast, L-DE-71 showed no significant difference, representing reduced recognition memory. Recognition Index (RI) analysis was not able to detect a memory deficit since mean values for all groups were not significantly different from the hypothetical value of 0.65 (one-sample t-test) as reported for WT (Tanimizu et al., 2017)) (**Fig. 3b**). Interestingly, H-DE-71 males had an apparently *lower* mean RI score indicating *stronger* memory (*P*=.08). In a separate cohort we examined investigation time with a new novel stimulus mouse on Day 2 to determine whether the reduction of investigation time on Day 2 is specific to social memory formation or simply disinterest. In **Figure 3c**, no significant reduction of investigation time was noted as expected for VEH/CON and H-DE-71, suggesting that the deficit in recognizing familiar mice in Fig. 3a was specific to social recognition memory formation. In contrast, the L-DE-71 group exhibited a significant reduction in investigation time of a different novel mouse on Day 2 (*P*<.05), indicating deficient social memory. **Figure 3 d** shows that RI scores for novel Day 2 memory were no different from the hypothetical value of “1” which would indicate disinterest (Tanimizu et al., 2017). In summary, these results indicate that developmental exposure to DE-71 at 0.1 mg/kg/d but not 0.4 mg/kg has mild effects on long-term social recognition memory.

**Fig. 3.**
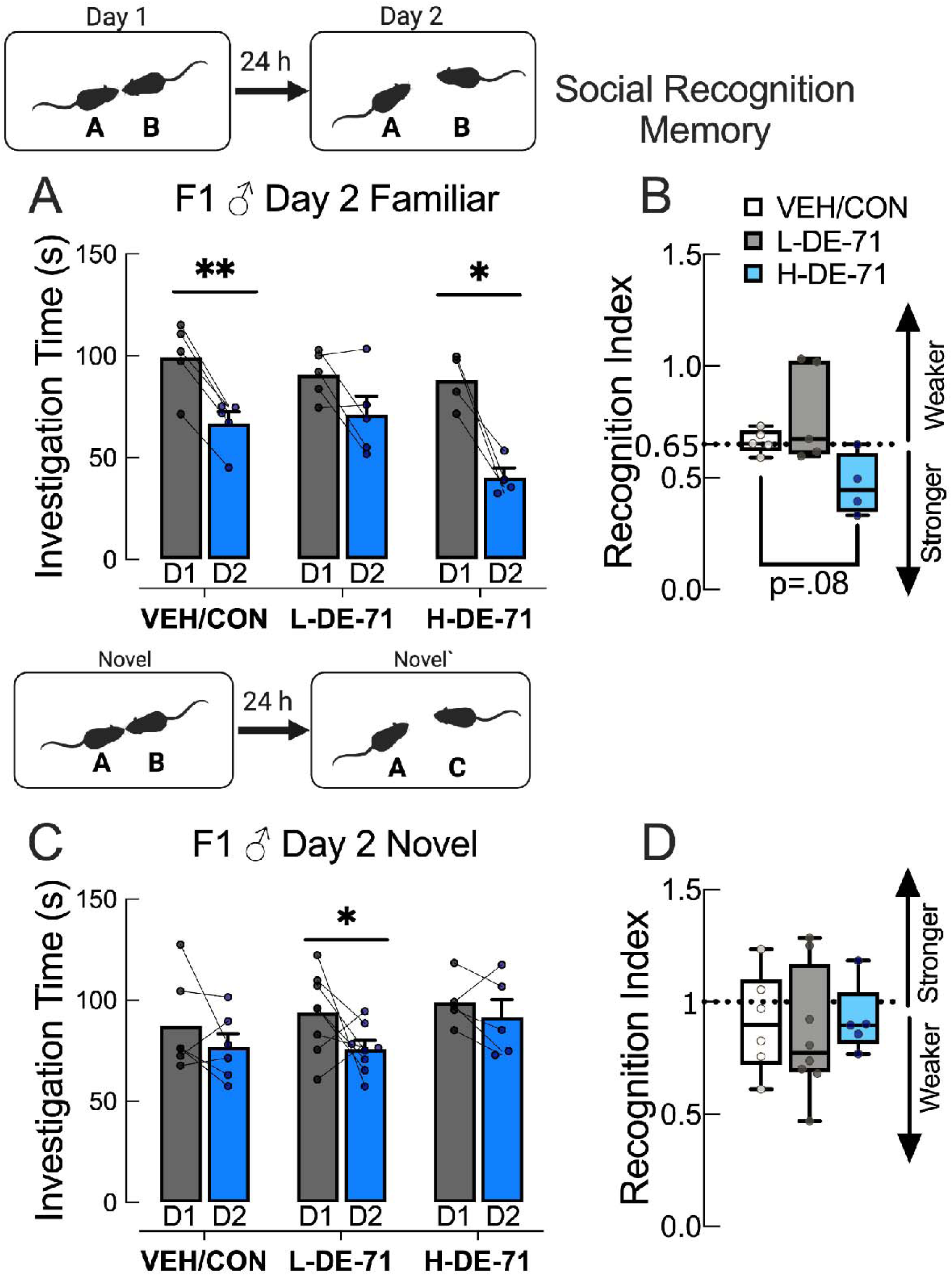
Exposure to L-DE-71 compromises long-term social recognition memory (SRM) in adult male mice. **a** When using the same novel stimulus mice on Day 2 (now familiar) the VEH/CON and H-DE-71 test mice (A) displayed a significant reduction in investigation time after the 24 h retention period on Day 2 relative to Day 1, indicating normal SRM. In contrast, L-DE-71 showed no significant difference, representing reduced SRM. **b** Corresponding Recognition Index (RI) scores for familiar were no different from 0.65, the hypothetical value representing SRM (one-sample t-test). **c** When presenting a novel mouse on Day 2 (C, Novel’), VEH/CON and H-DE-71 F1 mice showed equal preference for Novel’, confirming the face validity of the SRM construct. However, L-DE-71 mice displayed differential investigation of the second novel stimuli (Novel vs Novel’). **d** Mean RI scores for the novel were not less than the hypothetical value of 1.0, representing normal SRM in this paradigm (one-sample t-test). **\***compared to Day 1, \**P*<.05, \*\**P*<.01. *n*, 4-5 litters/group (a); 5-8 litters/group (c). A, test mouse; B, Novel; C, Novel’; D, day

### DE-71 compromised short-term novel object recognition memory

Having found that DE-71 exposure produces significant impairment in the SNP and SRM tests, we tested the hypothesis that DE-71 exposure also interferes with non-social recognition memory. Using a novel object recognition memory test (NORT), **Figure 4a** shows that L- and H-DE-71 males did not preferentially explore a novel object after a 30 min retention of familiar object on Day 1 testing, as did the VEH/CON (*P*<.05), indicating that DE-71 alters short-term memory for previously encountered objects. This was corroborated by discrimination index scores (DI) that were greater than 0 (no preference) only for VEH/CON (**Fig. 4b**, *P*<.05, one-sample t-test). Additionally, across group comparisons indicated that L-DE-71 had an apparently lower DI relative to VEH/CON (*P*=.07). On Day 2 mice were tested for long-term retention of object memory. The VEH/CON (*P*<.05) but not L-DE-71 nor H-DE-71 groups preferred novel over familiar objects (**Fig. 4e**). This was corroborated by the fact that DI scores were significantly different from 0 for VEH/CON (*P*<.01) but not L-DE-71 nor H-DE-71. These results suggest that DE-71, at both doses, interfered with short- and long-term object recognition memory. There were no effects of exposure on distance traveled (**Fig. 4c,d,g,h**), indicating that locomotor ability did not confound the NORT results.

**Fig. 4.**
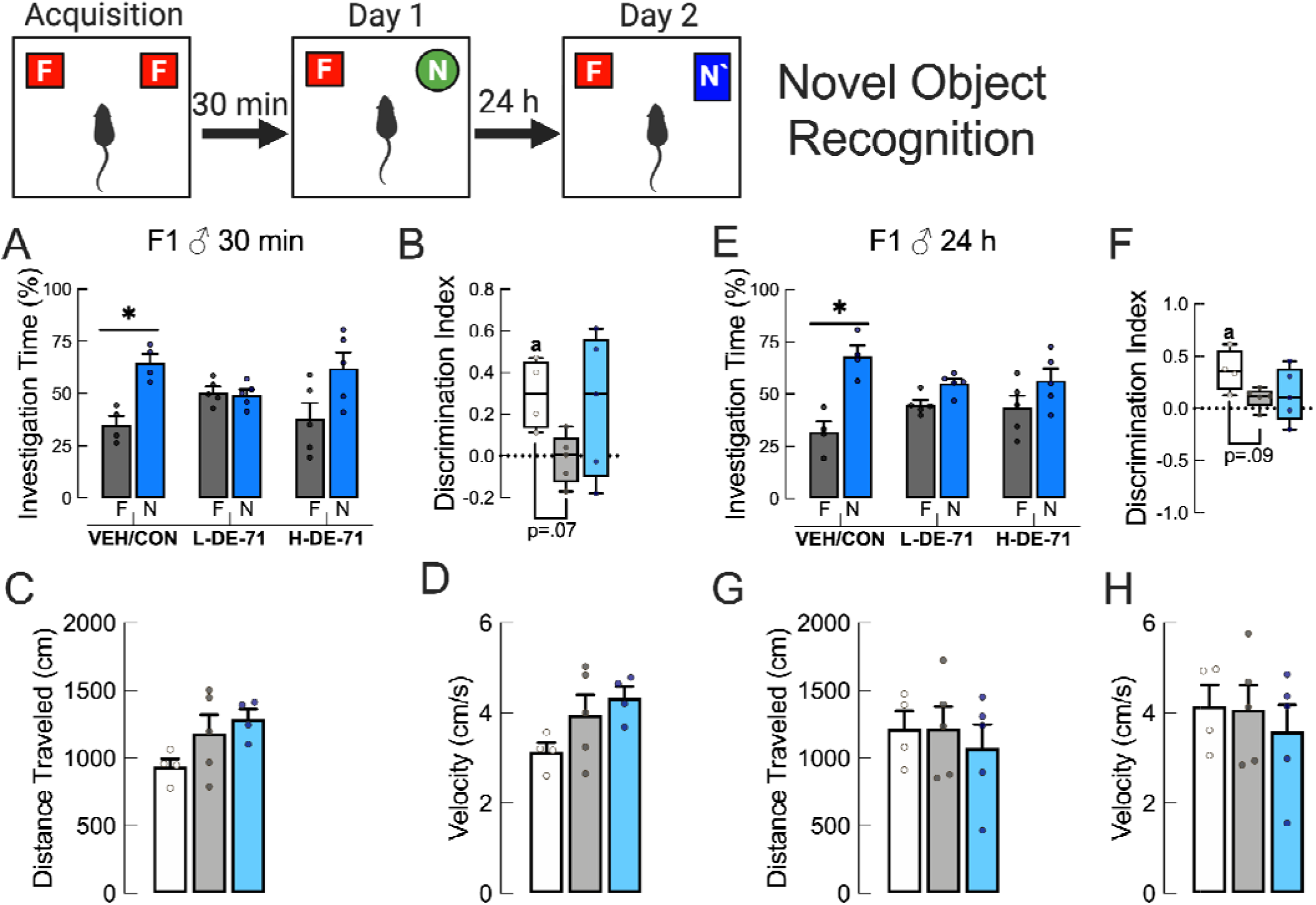
Perinatal exposure to L-DE-71 and H-DE-71 compromised short- and long-term novel object recognition memory in adult male mice. **a** Investigation time on short-term novel object recognition test. On the probe trial of Day 1 conducted 30 min after acquisition, exposed male offspring in the VEH/CON but not L-DE-71 or H-DE-71 groups show significantly greater time spent investigating the novel (N) vs familiar (F) objects. **b** Mean discrimination index scores for L- and H-DE-71 males were no different from 0 (no preference) in comparison to that of VEH/CON, which showed preference for the novel object (N). **e** A second probe trial conducted after a 24 h retention period measured mean investigation times of a second novel object (N’) indicated deficient object recognition for L- and H-DE-71 groups. **f** Mean discrimination index scores for *long-term* recognition memory were different from the hypothetical value of “0”, indicating no preference, for only VEH/CON. **c,d,g,h** distance traveled or velocity during the 5 min test. *compared to familiar object, \**P*<.05. **^a^**compared to hypothetical “0” score, ^a^*P*<.05, one-sample t-test. *n*, 4-5 litters/group. F, familiar object; N, novel object on Day 1; N’, novel object on Day 2

### Abnormal behavior produced by DE-71 exposure is not due to deficits in general olfactory processing

In order to examine if DE-71-induced deficits observed in SRT, SRMT, NORT were due to insufficient olfactory ability, we subjected male offspring to an olfactory preference test. **Figure 5a** shows that all mice, including those treated with DE-71, displayed preference for attractive vs aversive odorants, i.e., increased sniffing duration for peanut butter over water (*P*<.01, .001), butyric acid (*P*<.05, *P*<.01), and over vanilla (*P*<.05-.001).

**Fig. 5.**
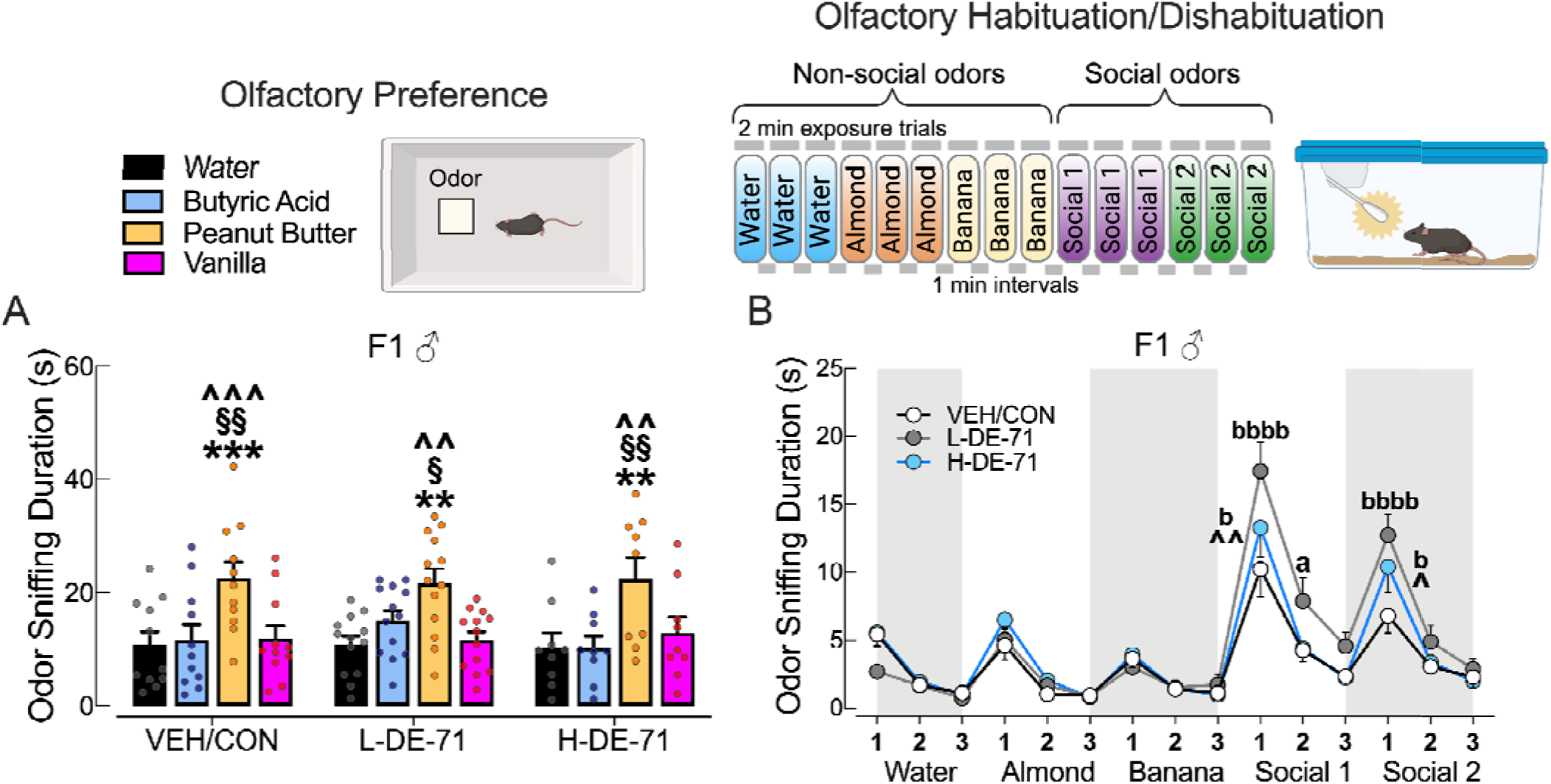
Perinatal exposure to DE-71 does not alter general olfaction but disrupts olfactory discrimination of social odors in adult male offspring. **a** DE-71-exposed male offspring showed normal olfactory preference. **b** Sniffing time on olfactory habituation/dishabituation test showed less habituation to social odor 1 in L-DE-71 vs VEH/CON. Both L-DE-71 and H-DE-71 showed abnormally reduced dishabituation from banana to social odor 1 and social odor 1 to 2. *Peanut butter compared to water (W), \**P*<.05, \*\*\**P*<.0001; §peanut butter compared to butyric acid (B), §*P*<.05, §§*P*<.01; ^peanut butter compared to vanilla (V), ^^*P*<.01, ^^^*P*<.001; **^a^**statistical difference in habituation to social odor 1 vs VEH/CON, ^aa^*P*<.01; **^b^**statistical difference in dishabituation to previous odor vs VEH/CON, ^b^*P*<.05, ^bbbb^*P*<.0001. Statistical results are summarized in Table 2. *n*, 9-13 litters/group (a), *n*, 16-18 subjects/group (b)

### DE-71 altered olfactory discrimination of social odors

We used an olfactory habituation/dishabituation test to measure the ability to discriminate non-social from social odors. All groups displayed olfactory habituation to all non-social odors and social odors (except L-DE-71 did not habituate to non-social odor 2-banana) as indicated by the decline in time spent sniffing odorant at trial 3 **(Fig. 5b**). Table 2 indicates the statistical results of the habituation/dishabituation test. When compared to VEH/CON group both L- and H-DE-71 groups displayed exaggerated dishabituation from non-social 2 to social odor 1 (*P*<.0001 and *P*<.05, respectively) and from social odor 1 to social odor 2 (*P*<.0001 and *P*<.05, respectively). Moreover, L-DE-71 showed deficient habituation to social odor 1 (*P*<.05). These results suggest that DE-71 produces reduced olfactory discrimination (hyposmia), especially of social odors, which requires processing via medial olfactory epithelium (MOE) and vomeronasal organ (VNO) (Huckins et al., 2013).

### DE-71 exposure does not promote anxiety nor depressive-like behavior

Male offspring were evaluated for anxiety using the EPM test (**Supplementary Figure 4**). Percent time spent in closed arms relative to open arms was significantly greater in all exposure groups (*P*<.0001). There was no effect of exposure on the number of open arm nor total arm entries. Using a forced swim test, that measures depressive-like behavior as time spent immobile, we found no significant group effects (**Supplementary Figure 4**).

### Effects of DE-71 exposure on Suok test

Using Suok, we measured locomotion, exploratory behavior, sensorimotor coordination and anxiety. There were no group effects detected. (**Supplementary Figure 5**).

### Early-life PBDE exposure does not alter locomotion on the Open Field test

The open field test informs about locomotion, habituation to novelty and anxiety. H-DE-71 groups showed hyperactivity, relative to VEH/CON, at PND 100, but not at PND 30 (**Fig. 6 a, d**). However, this occurred only during the first 15 min of the test (**Fig. 6 d**, *P*<.05, .01). These differences were taken into account when interpreting data on NORT and other tests. Time-dependent reduction in exploratory activity (habituation) was defined as reduced distance traveled (after 50 min) relative to the start of the test. All groups tested at PND 30 and 100 habituated to the arena with the exception of L-DE-71 males at PND 30 (**Fig. 6 a**, *P*<.05-.001).

**Fig. 6.**
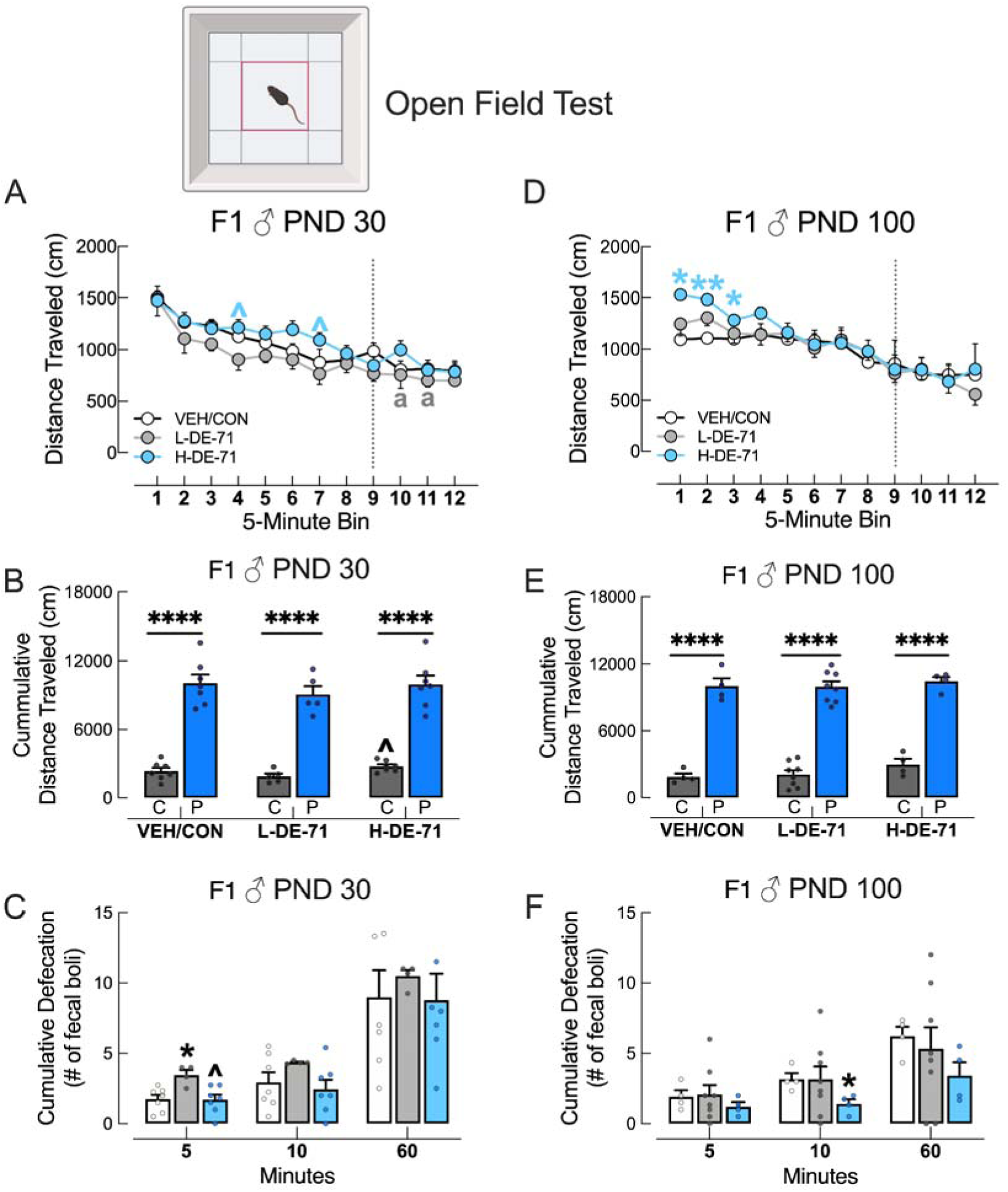
Effects of early-life PBDE exposure on the open field test. **a, d** Distance traveled in the open field arena indicated less habituation in L-DE-71 at PND 30 and early hyperactivity in H-DE-71 at PND 110. **b, e** Exploration time in center was less than in periphery for all groups, suggesting no exposure effects on anxiety. **f** Another measure of anxiety, number of fecal boli, indicated increased emotional reactivity in the L-DE-71 and H-DE-71 relative to VEH/CON. *compared to center (b,e) or VEH/CON (c,f), \**P*<.05, \*\*\*\**P*<.0001. ^compared to corresponding L-DE-71, ^*P*<.05. ^a^values at time bins after dashed line were not different from initial; Time bins lacking an “a” indicate they were significantly different, *P*<.01-.0001. *n*, 4-7 litters/group (a-c); 4-8 litters/group. C, center zone; P, periphery zone

**Fig. 7.**
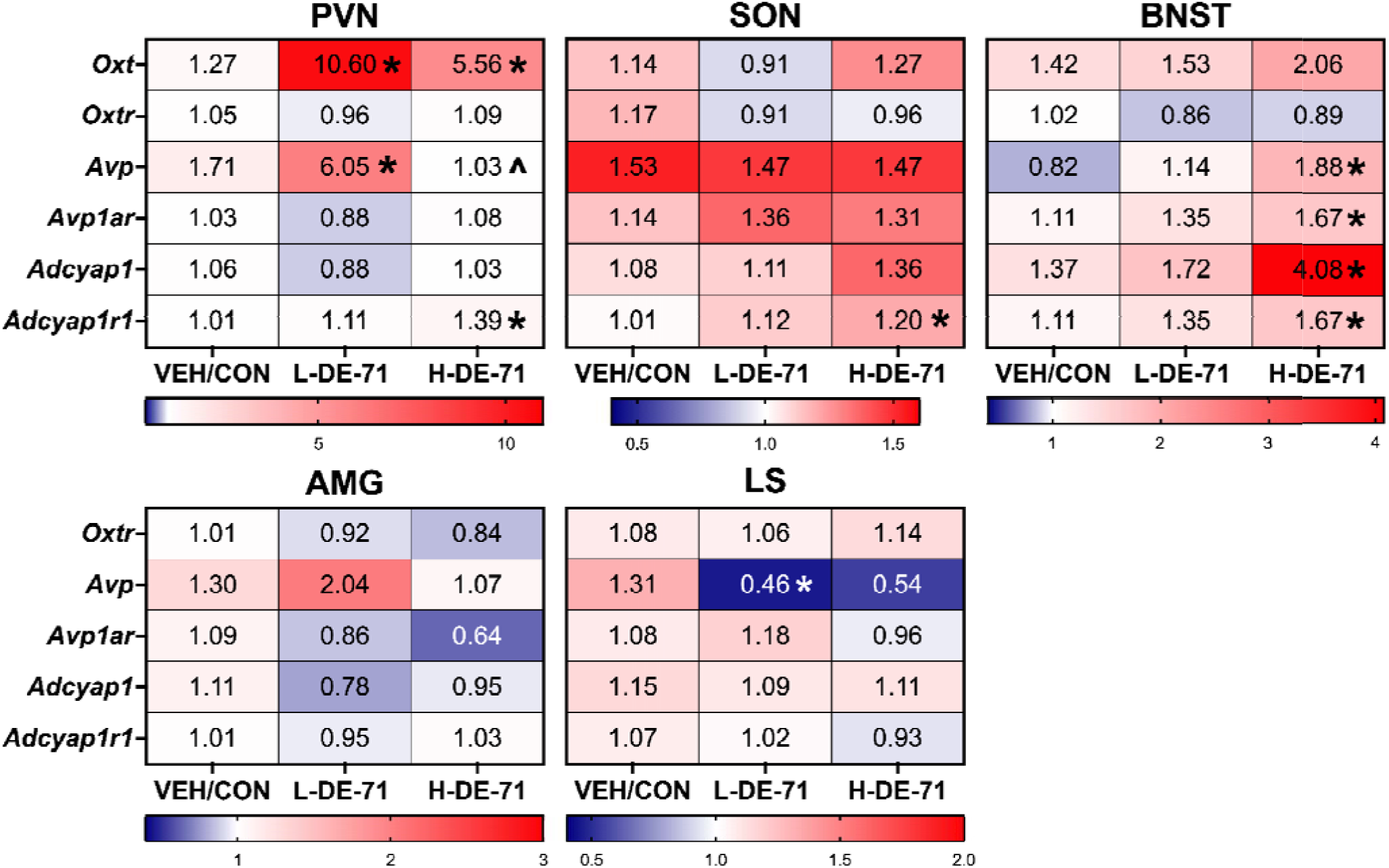
DE-71 exposure alters mRNA expression of social neuropeptide markers in select regions of the social neural network in male offspring. Heatmap representation (blue— downregulated; red—upregulated) of RT-qPCR analysis displaying the mean fold-change of each gene studied by brain region in selected SNN brain regions harvested at PND 110. *compared to VEH/CON, \**P*<.05-.01. ^compared to L-DE-71, ^*P*<.05. PVN, paraventricular nucleus; SON, supraoptic nucleus; BNST, bed nucleus of the stria terminalis; AMG, amygdala; LS, lateral septum; *n*, 3-14 samples/group

Exploration in the periphery vs center zones was greater for all exposure groups at both ages as expected for the brightly lit conditions (**Fig. 6 b,e**, *P*<.0001). **Figure 6 b** shows that total distance traveled in the center was greater for H-DE-71 versus L-DE-71 (*P*<.05). Another measure of anxiety, number of fecal boli when first exposed to open field arena at 5 min, indicated increased emotional reactivity in PND 30 L-DE-71 relative to VEH/CON (**Fig. 6 c,** *P*<.05). In contrast, at PND 100 H-DE-71 had fewer fecal boli (*P*<.05) (**Fig 6 f**).

### DE-71 exposure alters mRNA expression of social neuropeptide markers in social neural network regions

To correlate the behavioral findings with deregulated gene markers of neuropeptide systems, that are key mediators of complex social behavior, we measured relative mRNA expression for vasopressin (*Avp*), oxytocin (*Oxt*), PACAP (*Adcyap1)* and their receptors. mRNA was obtained from micropunches of discrete brain nuclei involved in social behavior: amygdala (AMG), bed nucleus of the stria terminalis (BNST), lateral septum (LS), PVN and SON and processed using RT-qPCR. **Figure 8** shows that *Oxt* mRNA transcripts were increased in L-DE-71 PVN (P<.01) and H-DE-71 PVN (*P*<.05). *Avp* was downregulated in L-DE-71 LS (*P*<.05) and apparently downregulated in H-DE-71 LS (*P*=.07) but upregulated in L-DE-71 PVN (*P*<.01) and H-DE-71 BNST (*P*<.05). For the PACAP gene, *Adcyap1,* mRNA transcripts were upregulated in H-DE-71 BNST *(P*<.05). With regard to receptors, *Oxtr* levels were *apparently* decreased in H-DE-71 AMG vs VEH/CON when using t-test (*P=*.08) but not when using ANOVA. For *Avp1ar,* H-DE-71 BNST levels were upregulated (*P*<.05). For the PACAP receptor gene, *Adcyap1r1,* mRNA transcripts were upregulated in H-DE-71 PVN (*P*<.01), SON (*P*<.05), BNST (*P*<.05).

**Fig. 8.**
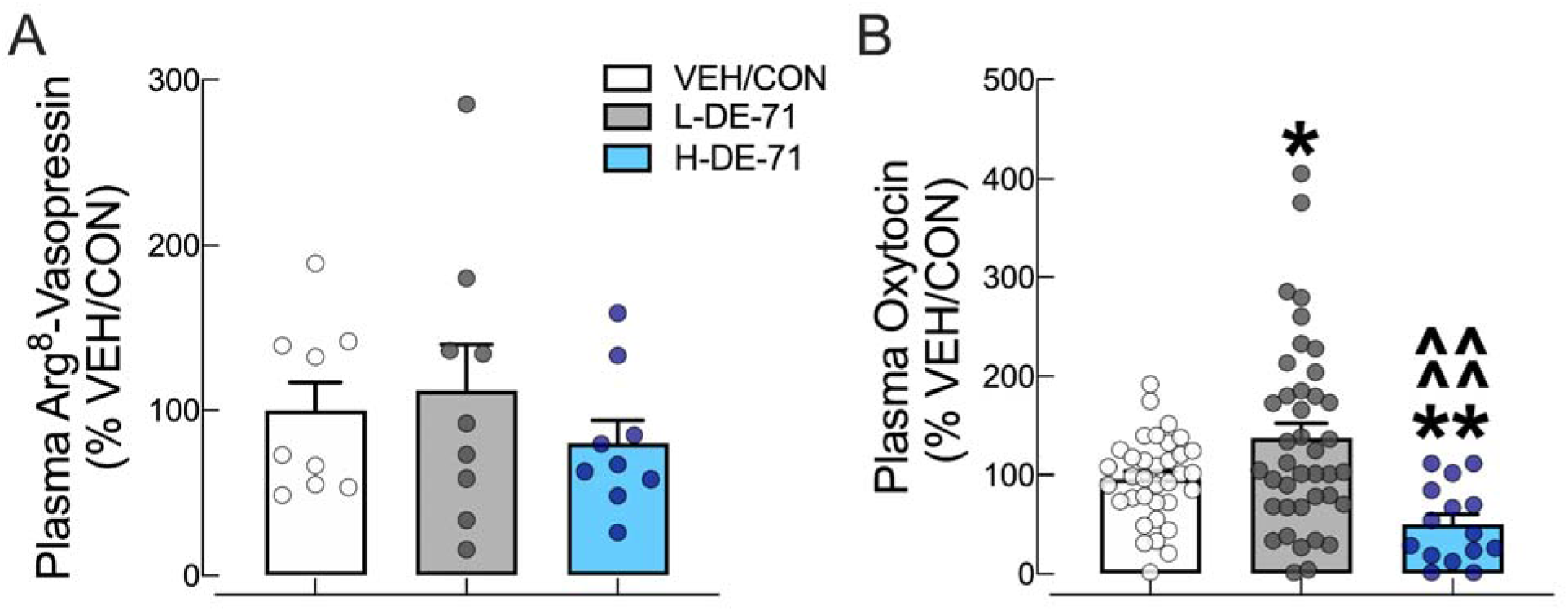
Perinatal exposure to DE-71 produced dose specific responses to plasma OXT level in adult F1 male offspring. **a** Plasma Arg-8 vasopressin measured using EIA using blood taken from adult male offspring at sacrifice. **b** Oxytocin levels showed dose-dependent increase or decrease in L-DE-71 and H-DE-71 respectively relative to VEH/CON. *compared to VEH/CON, *P<.01, \*\**P*<.01; ^compared to L-DE-71, ^^^^*P*<.0001. *n*, 9 subjects/group (a); *n*, 15-40 subjects/group (b).

### DE-71 Produced Dose Depended Changes in Adult Plasma Levels of Oxytocin

We next measured plasma OXT and Arg8-vasopressin concentrations to examine their association with social behavior phenotypes. **Figure 9a** shows that plasma AVP levels were not changed with DE-71 exposure. In contrast, plasma OXT levels increased in L-DE-71 (P<.05) decreased in H-DE-71 males (*P*<.01), indicating a dose-specific effect (**Fig. 8 b**; *P*<.0001).

## Discussion

Mounting evidence links early-life exposure to environmental toxicants with an increased risk of neurodevelopmental disorders (NDDs), however, a causal link has not yet been established and the specific ways in which such exposures intersect with biological pathways relevant to ASD is not completely understood. Our study addresses this gap by demonstrating that developmental exposure to the penta-PBDE mixture DE-71 induces long-lasting impairments in ASD-relevant outcomes such as social recognition, repetitive behavior and social odor discrimination in offspring. These behavioral alterations were accompanied by changes in plasma oxytocin and gene markers of OXT and AVP signaling in the social neural network. By modeling human-relevant exposure through the full complement of congeners detected in breast milk, and using the litter as the unit of analysis to minimize bias and inter-individual variability, this study provides additional evidence for the neurobehavioral impact of early-life exposure to PBDEs. Importantly, this work extends our previous findings in females to include male offspring, revealing sex-specific effects of PBDE exposure on social behaviors and neuroendocrine pathways. Collectively our findings underscore the heightened vulnerability of the developing brain to environmental xenobiotics during critical windows of neurodevelopment.

The most salient finding of this study is that perinatal transfer of DE-71 from exposed mothers produces behavioral phenotypes in their male offspring that mirror the two core symptom domains of ASD diagnosis: deficits in social reciprocity and communication and repetitive or stereotyped behaviors (American Psychiatric Association, 2013). Specifically, both doses of DE-71 significantly reduced mean scores on social novelty preference (short-term social recognition) and/or social recognition memory tests, but sociability was not affected. Notably, the impairment in social recognition memory seen in juveniles persisted into adulthood, indicating an enduring effect of perinatal exposure. In other words, PBDEs selectively disrupt specific domains of social cognition. They impair performance on the “*knowledge of self and others*” domain (short- and long-term social recognition) but not “*social motivation*” (sociability) (Bicks et al., 2015). In humans, the ability to recognize other individuals is essential for the formation of any type of enduring social relationship. Notably, early social recognition is disrupted in children with ASD and performance on a facial recognition task has been shown to predict ASD symptom severity (Dawson et al., 2012). Together, these findings provide evidence that PBDEs in early life disrupt normal social behavior translatable to ASD, with enduring effects into adulthood.

In addition, the same DE-71-exposed mice displayed significantly exaggerated repetitive behavior on a marble burying test indicative of disruption across a second core domain of ASD. These behavior defects are not likely due to either an overall movement deficit or heightened anxiety, both of which can also signal neurodevelopmental disruption. While we observed hyperactivity during the early part of OFT in H-DE-71-exposed mice, it was only seen at PND 110, while SRM deficits were seen in both L-DE-71 and H-DE-71 males at PND 30. Others have found hyperactivity after acute exposure to 0.8 mg/kg BDE-99 at PND 10, suggesting that some congeners in DE-71 may be the active contributors (Viberg et al., 2004); (Costa & Giordano, 2007). We observed co-occurring disruptions on other constructs that are involved in SRM ability, namely olfaction (hypo- and hyper-sensitivity in odor detection) and hippocampal memory (Raam et al., 2017; Scheier et al., 2024; Tanimizu et al., 2017), that are related to clinical markers and co-morbidities in ASD (Ashwin et al., 2014; Khachadourian et al., 2023). For example, SRM disruption was linked to exaggerated olfactory dishabituation of social odors shown by L- and H-DE-71 exposed male mice. SRM deficits in AVP1AR null mice can occur in the absence of any impairments in spatial and non-social olfactory learning and memory tasks (Bielsky et al., 2004). PBDE exposure also produced reprogramming of hippocampal and related neural circuits orchestrating other related memory functions, i.e., object recognition memory. As with DE-71-exposed females studied previously (Kozlova et al., 2021), reduced mean scores on NORT indicated additional memory impairment in exposed male mice. In analogy, children with ASD often suffer from behavioral and intellectual co-morbidities beyond core symptoms such as ADHD and intellectual disability (Lord et al., 2018). The social behavior deficits are similarly reported for other mouse models of autism, although information on the corresponding status of locomotor activity, olfaction, and hippocampus-mediated memory in these models is scarce. Our findings provide a broader characterization of the abnormal phenotypes that are relevant to ASD core symptoms and co-morbidities produced by early-life exposure to environmentally and humanly relevant EDCs such as PBDEs.

Human epidemiological studies examining the association between body burden of PBDEs and social or ASD-related outcomes have yielded mixed findings. Some report associations between postnatal BDE-47 exposure and reduced social competence in children (Braun et al., 2014; Ding et al., 2015; Gascon et al., 2011; Hartley et al., 2022; Song et al., 2023), while Hamra and others (2021) observed that each unit increase in PBDE burden was linked to a 1.41-fold higher odds of ASD risk (Hamra et al., 2021). Other studies, however, found weaker, inverse, or null associations (Hertz-Picciotto et al., 2011; Lyall et al., 2017; Vuong et al., 2018). Experimental ASD models mirror a comparable lack of consensus, likely due to type of PBDE congeners, dose, exposure timing, sex, and species. Evidence from kestrels exposed to DE-71 aligns with our results, demonstrating abnormal pair-bonding and courtship behaviors (Fernie et al., 2005). Other reports on BDE-47 describe reduced sociability at (0.2 mg/kg) <u>(Kim et al.</u> <u>2015)</u>, but no effect on recognition ability unless combined (at 0.03 mg/kg) with *Mecp2* deficiency at <u>(Woods et al. 2012)</u>, despite reduced exploratory activity when high doses are used (50 mg/kg) (Z. Li et al., 2021). Studies of low-dose BDE-209 (0.12 ng/mouse/day, s.c.) also found no social recognition deficits in male mouse offspring (Chen et al., 2019). We speculate that neither BDE-99 nor BDE-209 alone is sufficient to induce deficits in social recognition or social memory even used in a range of doeses. However, early postnatal exposure to these congeners has been shown to cause cognitive impairment and hyperactivity (Branchi et al., 2002, 2003; Costa & Giordano, 2007; Eriksson et al., 2001; Viberg et al., 2004). Thus, it is more likely that multiple PBDE congeners act synergistically and/or additively to produce the disrupted social phenotypes. This interpretation is reinforced by findings obtained with other flame retardants: perinatal exposure to the EDC mixture, NeuroMix, Firemaster 550 or its individual components, disrupts social recognition and partner preference in a sex-dependent manner in rats and voles (Gillera et al., 2020; Gore et al., 2022; Witchey et al., 2020). Similarly, exposure to PCBs has been linked to altered sociosexual preference and social investigation (Hernandez Scudder et al., 2020; Reilly et al., 2015). Research results from these combined studies warns of the ability of developmental exposure to PBDEs and other EDCs to reprogram social brain and general memory circuits leading to lasting social cognition abnormalities.

Previous work from our group has shown that both in vitro and developmental exposure to PBDEs (or PCBs) can interfere with neuroendocrine regulation of the prosocial neuropeptide Arg8-vasopressin (AVP) with physiological stimulation (Coburn et al., 2005, 2007; Mucio-Ramírez et al., 2017). These findings suggest that the PBDE-induced impairments in SNP and SRM seen in the current study may stem from disrupted OXT signaling and/or interference with the associated AVP system. OXT has pro-social as well as other effects (Ring et al., 2006) and a causal link between OXT disruption and ASD has been suspected but has not been definitively established. For example, in unexposed animals blocking the OXT system by gene deletion (OXT, CD38, OXTR) (Ferguson et al., 2000; Jin et al., 2007; Lee et al., 2008; Sala et al., 2011; Takayanagi et al., 2005) or by pharmacological blockade in AMG will impair SRM (Ferguson et al., 2001). Our PCR results reveal sex-specific deregulation of *Oxt, Avp,* and their receptors in key SNN hubs that may provide mechanistic insight into behavioral deficits produced by both doses of DE-71. Specifically, *Oxt* mRNA levels were elevated in male PVN and reduced plasma OXT, in contrast to the previously reported reduction in female BNST and SON concomitant with *Oxtr* upregulation in several regions of the female SNN (BNST, AMG, PVN) (Kozlova et al., 2021). Reilly and others (2022) have reported elevated *Oxt* in male PVN after developmental exposure to Aroclor1221, suggesting this as a common target of EDC neurotoxicity (Reilly et al., 2022). Manipulation of OXT signaling has been attempted in clinical trials to ameliorate autistic symptoms but with inconsistent results. For example, in adolescents and adults autistic individuals, OXT was found to improve social deficits, such as face recognition and social responsiveness (emotion recognition and interaction) related to ASD or other neurodevelopmental disorders (Anagnostou et al., 2012; Guastella et al., 2015; Peñagarikano, 2015). Also, intranasal OXT treatment can enhance social abilities in children with ASD, especially those with the lowest pretreatment OXT concentrations (Parker et al., 2017)

AVP also controls social processes that are critical for the formation and maintenance of social relationships: social recognition, social communication and aggression by modulating the interpretation of sensory information, influencing decision making and by triggering complex motor outputs (Albers, 2012). Regulation of social behavior by AVP may be more predominant in males via AVP1a receptors in BNST and LS (Rigney et al., 2024)(Bluthé & Dantzer, 1990). Here, we found that DE-71 exposure deregulated the male AVP system: *Avp* was upregulated in PVN (L-DE-71) and BNST (H-DE-71) and downregulated in LS (L-DE-71) concomitant with upregulation of *Avp1ar* in BNST (H-DE-71). Although DE-71-exposed females also show upregulation of *Avp1ar* in BNST (and downregulation in SON), V1aR antagonists are only able to block social recognition in males (Bluthé & Dantzer, 1990). Because the promoter regions of OXT (Mamrut et al., 2013) and AVP genes (Auger et al., 2011) are sensitive to epigenetic modification, developmental BDE-47 exposure may alter their regulation through global DNA methylation changes (Poston & Saha, 2019; Woods et al., 2012). Yet another factor influencing sex differences may be due to the estrogenic, anti-estrogenic and androgenic activity of some PBDE congeners since these genes are regulated by reproductive hormones (Ren & Guo, 2013). Early life exposure to various EDCs (PCBs, Firemaster 500, Bisphenol A) have also been demonstrated to modify the hypothalamic OXT and AVP neurochemical systems in a sex-, region-, and treatment-specific manner. EDCs such as Firemaster 550 (Gillera et al., 2021), PCBs including Aroclor 1221 and 1254 (Coburn et al., 2005; Gore et al., 2023; Reilly et al., 2022), and Bisphenol-A (Witchey et al., 2019) disrupt OXT and/or AVP systems, effects that parallel their impacts on social behavior (Gillera et al., 2020; Gore et al., 2019, 2023; Witchey et al., 2020).

PBDEs have been mostly studied in regard to their endocrine disruption of thyroid and reproductive systems (Costa & Giordano, 2007; Zhao et al., 2015). While the mechanisms underlying the abnormal neurobehavioral phenotypes are unclear, the pleiotropic effects of thyroid hormone (TH) in mediating key neurodevelopmental events is likely contributing (Costa & Giordano, 2007). Animals rendered hypothyroid exhibit behavioral alterations that include hypo-and hyperactivity, learning and memory impairment, and altered anxiety and olfaction (Cheng et al., 2025; Umezu et al., 2019; Vancamp et al., 2022; Vasilopoulou et al., 2016). Few studies have mechanistically linked PBDE-induced TH disruption to neurobehavioral alterations. In particular, De Miranda et al. (2016) found that, in female DE-71 exposed rats, reference memory impairment was improved after thyroxine (T4) supplementation (de-Miranda et al., 2016). With regards to social behavior, we recently found that maternal T4 supplementation improved TH status as well as social recognition memory deficits in DE-71 exposed mice (Kozlova et al., 2025). Based on these findings, we speculate that PBDE-induced disruption of TH signaling may underlie PBDE-mediated abnormal cognition and social phenotypes as speculated previously (Currás-Collazo, 2011; Kodavanti & Curras-Collazo, 2010; Kozlova et al., 2025; Martin et al., 2025).

Using GC/MS we confirmed penetration of the major BDE congeners in DE-71 in juvenile brain of male offspring at PND 30, with the sum concentration (<150 ng/g ww or 0.15 ppm) lower than in our previously reported female brain samples at PND 15 (∼250 ng/g ww) likely due to separation from the maternal PBDE source in males by PND 30. Serum ∑PBDE were measured to reach 482 ng/g l.w. in California toddlers (Fischer et al., 2006). Using a divisor factor of .097 to convert w.w. to l.w. (unpublished observations), we estimate our mean brain ∑PBDE in H- and L-DE-71 males at PND 30 to be similar, i.e., 1.0- or 3.1-fold greater, respectively, suggesting our mouse model of maternal PBDE transfer is translational. As with females, lower but significant ppm levels of ∑PBDEs still remained into adulthood in male brains at PND 110. The congener profile in PND 30 brain samples was dominated by BDE-47, - 99, −139, −100, −153, and −154, which accounted for 97% of the mean ∑PBDEs. These congeners also comprise the majority of congeners (97%) in DE-71 (Kodavanti et al., 2010). In general, this congener composition mimics that found in toddler serum and breastmilk, i.e. BDE-47, −99, - 100, −153 and −154 (Cowell et al., 2019; Fischer et al., 2006; Rose et al., 2010; Schecter et al., 2005). Additional congeners detected in offspring brains, i.e., BDE-17, 49, 138, 139, 140, 183 and 184, have also been detected in human serum and/or breastmilk, signifying that the exposures modeled in this study reflect environmentally relevant levels and congener profiles in humans (Chao et al., 2010). Postmortem brain samples from 4-71 year-olds born 1940 to 2000 have also been shown to contain BDE-26, −47, −99, −100, −153, −154 and −183 at ppb levels, but levels are not higher in autistics vs non-autistic controls although autistics contained reduced BDE-153 (Mitchell et al., 2012). This congener profile may be attributed to several possible commercial formulations to which humans are exposed. When compared to females, male juvenile brains contained similar BDEs, but in adulthood, males showed relatively more remaining congeners (BDE 28, 47, 100, and 153) as compared to adult female brain samples in which only BDE-153 was detected (Kozlova et al., 2021). In particular, BDE-47 has been positively associated with autistic features and is the most prevalent PBDE congener in breastmilk (Braun et al., 2014; Ding et al., 2015; Gascon et al., 2011; Hartley et al., 2022; Song et al., 2023). Notably, a recent review indicates that some of these congeners are still detected in diets sampled worldwide (Gao et al., 2024).

Sex-specific differences are evident in the effects of developmental PBDE exposure when comparing our present findings using males with previous findings in females (Kozlova et al., 2021). In our model, DE-71-exposed males exhibited less pronounced deficits in long-term social recognition memory compared to exposed females. In addition, the slightly greater dose, H-DE-71, reduced SNP in males while L-DE-71 was the only dose that produced a phenotype in females. Repetitive behaviors are equally exaggerated in L-DE-71 of both sexes while altered outcomes of social odor discrimination were sex-specific, i.e., decreased in L-DE-71 females and exaggerated in L-DE-71 males. Deregulation of hypothalamic pro-social gene markers in males depicts a different profile than in females. Specifically, *Oxt* was regulated oppositely in females (down) and males (up). Sex-specific gene deregulation was also noted for *Avp* in BNST, being downregulated in females but upregulated in males. Only males showed *Avp* alteration in lateral septum and *Oxtr* upregulation was only seen in females. Both sexes showed upregulated *Avp1ar* in BNST. The significance of these changes may be related to sexually dimorphic circuitry controlling social recognition ability (Albers, 2012). Sex differences observed in our study may reflect the interaction of PBDEs with sexually dimorphic neuroendocrine systems, including OXT and AVP pathways that play critical roles in social cognition, but that are differentially regulated in males and females (Dumais & Veenema, 2016).

### Conclusions

Applying established behavioral assays to evaluate ASD-relevant behaviors, we observed that DE-71 exposure specifically impaired short- and long-term social recognition memory, increased repetitive behaviors, and disrupted olfactory discrimination of social odors. Other behavioral domains including anxiety, depression, sensorimotor integration, hyperactivity or locomotion were largely unaffected, indicating that DE-71’s effects are targeted to ASD-relevant phenotypes rather than general neurological dysfunction. In our environmental model of ASD, we concomitantly observed alterations in prosocial neurotransmitters/neurohormones that play key roles in the social neural network. The OXT and AVP systems, in particular, are strongly implicated in social cognition and disruptions in these pathways have been associated with socio-behavioral impairments resembling the core features of autism (Aspé-Sánchez et al., 2015). Because of their central role, these neurochemical systems are currently being explored as promising therapeutic targets for ASD (Cataldo et al., 2018; László et al., 2023; Meyer-Lindenberg et al., 2011). Although past rodent and human studies examining an association between EDCs and ASD risk have yielded mixed results, our findings strengthen the evidence that xenobiotic toxicants may pose neurodevelopmental risk in males. Still, only a limited number of studies have directly investigated PBDE exposure in relation to autism-like phenotypes, and outcomes remain inconsistent, highlighting the pressing need for more detailed epidemiological and experimental research on persistent organic pollutants (POPs) and ASD vulnerability.

These findings indicate that early-life stages, including fetal and postnatal periods, represent a critical window of heightened vulnerability to the neurobehavioral effects of PBDEs. Previous studies have similarly reported that developing organisms are particularly sensitive to organohalogen-induced neurobehavioral disruptions due to the limited detoxification capacity of the developing brain and the disruption of key processes governing growth, differentiation, and neural circuit formation (Costa & Giordano, 2007). Our results show that deficient social recognition memory and exaggerated marble burying were still abnormal in adulthood. In the context of ASD, the developmental origins of adult disease framework, can be used to attribute long-lasting social and repetitive behavior and cognitive deficiencies to perinatal DE-71 exposure.

### Limitations of the Study

Behavioral analyses were mainly analyzed by litter as the unit of statistical analysis. However, due to limited resources, some tests used individuals. Brain BDE congener levels were quantified with two independent MS methods (GC/ECNI-MS, HRGC/HRMS), confirming that the decline by PND 110 reflected elimination rather than analytical error. PCR analysis showed sex-specific effects of PBDEs on prosocial neuropeptide genes and receptors in select social brain regions. Low cell density in areas such as the amygdala and LS reduced RNA yield and increased variability for low-abundance genes. To mitigate this, we optimized primer design per MIQE guidelines, though technical limitations remain.

Our focus has been on social behavior and HNS system however, it is known that exposure to PBDEs has been associated with a multitude of other health effects including liver injuries, TH disorders, neurotoxicity, genotoxicity and immune system disorders (Gao et al., 2024). In fact, we have reported that mice exposed to developmental PBDEs and directly exposed dams exhibit features of type 2 diabetes and metabolic syndrome, effects which may relate to the present findings (Kozlova, Chinthirla, et al., 2022; Kozlova, Denys, et al., 2022; Kozlova et al., 2020).

**Table 1.**
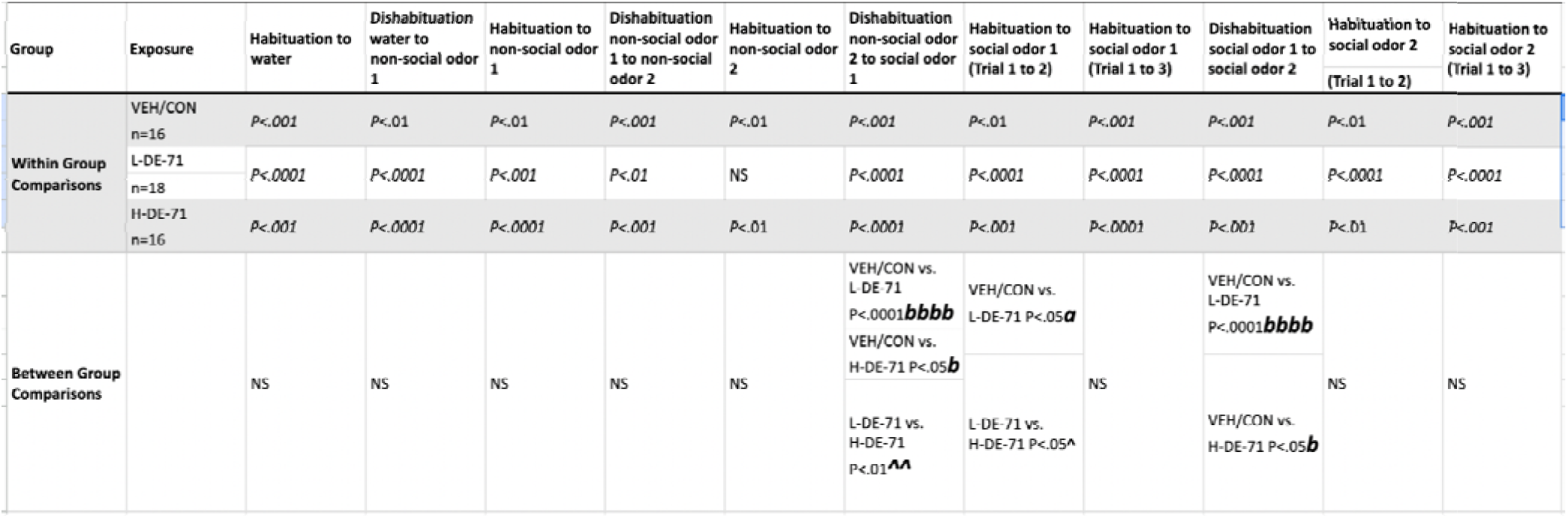
Statistical Results for the Olfactory Habituation/Dishabituation test Olfactory Habituation/Dishabituation test. Data and symbols are shown in Figure 5C,D. Non-social odor 1, almond; non-social odor 2, banana; social odors were obtained from dirty cages of sex-matched unfamiliar conspecifics ^a^habituation to previous trial of same odor vs VEH/CON, ^a^P<.05 ^b^dishabituation to previous odor vs VEH/CON, ^b^P<.05, ^bb^P<.01, ^bbbb^P<.0001 ^^^compared to L-DE-71, ^P<.05, ^^P<.01

## Acknowledgements

We thank Drs. M. Valdez, J. Valdez, H. Cherukury and P.R.S. Kodavanti for their contributions to this work. We acknowledge Drs. G. Hicks, M. Collin, D. Carter and C. Clark and H. Clark at UCR Institute for Integrative Genome Biology Genomics and Imaging Cores, and E. Grace (IDT) and M. Kuhn (NEB) for advice on PCR analysis. We thank UCR graduate and undergraduate students R. Bottom, H. Chen, J. Dansie, Z. Amador, A. Khan, K. Conner and D. Rohac for technical assistance. We thank Drs. K. Huffman (Dept. Psychology), W. Saltzman (Dept. Biology) for access to behavioral apparati and Dr. R. Calma (visiting scholar, Dept. Cell Biology & Neuroscience) for guidance on PCR methods. We are grateful to Drs. B. Wong (Noldus) and R. Hartman (Loma Linda University) for training and access to Ethovision. Dr. J. Porter and M. Colon at the Brain Behavioral Core (RR003050/MD007579), Ponce Health Sciences University, provided additional support on Ethovision data analysis. We thank A. Lam for help with editing the manuscript. We are grateful to Drs. I. Ethell, F. Sladek, K. Huffman, M. Riccomagno for gift of mice used as breeders and stimulus animals. Illustrations were created with BioRender.com

## Declarations

### Disclaimers

Research reported in this publication was supported by the National Institute Of Environmental Health Sciences of the National Institutes of Health under Award Number F31ES034304. The content is solely the responsibility of the authors and does not necessarily represent the official views of the National Institutes of Health.

The research described in this article has been reviewed by the Center for Public Health and Environmental Assessment, U.S. Environmental Protection Agency (EPA) and approved for publication. Approval does not signify that the contents necessarily reflect the views and policies of the agency nor does the mention of trade names of commercial products constitute endorsement or recommendation for use.

### Availability of Data and Material

Not applicable.

### Code Availability

Not applicable.

### CRediT authorship contribution statement

**Conceptualization:** E.V.K., M.C.C.-C.; **Methodology:** E.V.K., G.M.G., M.E.D., J.M.K., G.L., A.L.P., H.M.S., B.H., K.-W.S., M.C.C.; **Validation:** E.V.K., H.M.S., B.H., K.-W.S., M.C.C.; **Formal Analysis:** E.V.K., G.M.G., M.E.D., A.E.B., J.M.K., N.L., J.T., L.C., L.M.A., C.N.L., G.C., A.L.P., H.M.S., B.H., K.-W.S., M.C.C.-C.; **Investigation:** E.V.K., G.M.G., M.E.D., A.E.B., R.G., T.M., J.M.K., G.L., N.L., K.M.R., J.T., L.C., L.M.A., C.N.L., D.S.O., E.M., V.C., J.D.T., D.P., Y.K., B.D.C., M.B., S.K., A.L.P., B.H., M.C.C.-C.; **Writing – Original Draft:** E.V.K., M.E.D., A.E.B., M.C.C.-C.; **Writing – Review & Editing:** E.V.K., M.E.D., R.G., G.L., N.L., K.M.R., J.T., L.M.A., C.N.L., E.M., M.B., S.K., A.L.P., M.C.C.-C.; **Visualization:** E.V.K., G.M.G., A.E.B., J.M.K., M.C.C.-C.; **Resources:** E.V.K., G.M.G., M.E.D., J.M.K., H.M.S., K.-W.S., M.C.C.-C.; **Data Curation:** E.V.K., G.M.G., M.E.D., A.E.B., R.G., T.M., J.M.K., G.L., N.L., K.M.R., J.T., L.C., L.M.A., C.N.L., D.S.O., E.M., V.C., J.D.T., D.P., Y.K., B.D.C., M.B., S.K., G.C., A.L.P., H.M.S., B.H., K.-W.S., M.C.C.-C.; **Supervision:** E.V.K., G.M.G., M.E.D., H.M.S., B.H., K.-W.S., M.C.C.-C.; **Project Administration:** E.V.K., G.M.G., M.E.D., J.M.K., H.M.S., K.-W.S., M.C.C.-C.; **Funding Acquisition:** E.V.K., G.M.G., M.E.D., J.M.K., G.L., N.L., K.M.R., L.C., L.M.A., V.C., A.L.P., H.M.S., K.-W.S., M.C.C.-C..

### Ethics approval

Care and treatment of animals was performed in accordance with guidelines from and approved by the University of California, Riverside Institutional Animal Care and Use Committee (AUP# 5, 00170026 and 20200018).

### Consent to participate

Not applicable.

### Consent for publication

All authors reviewed and approved the final manuscript.

## Abbreviations

*ActB*: Beta Actin (reference gene)
ADHD: Attention Deficit Hyperactivity Disorder
*Adcyap1*: Adenylate Cyclase Activating Polypeptide 1 (PACAP)
*Adcyap1r1*: Adenylate Cyclase Activating Polypeptide 1 Receptor 1 (PAC1R)
AMG: Amygdala
ANOVA: Analysis of Variance
AVP: Arginine8-Vasopressin (*Avp*)
ASD: Autism Spectrum Disorder
*Avp1ar*: Arginine Vasopressin Receptor 1A
AUP: Animal Use Protocol
BDE: Brominated Diphenyl Ether (e.g., BDE-47, BDE-85, BDE-209)
BNST: Bed Nucleus of the Stria Terminalis
BFR: Brominated Flame Retardant
BORIS: Behavioral Observation Research Interactive Software
bw/d: Body Weight per Day
CHEM TUBE: Hydromatrix Chemical Absorbent (Agilent Technologies)
CON: Control
DCM: Dichloromethane
DE-71: Commercial PBDE mixture (technical grade)
ECNI: Electron Capture Negative Ionization
EDC: Endocrine Disrupting Chemical
EI: Electron Impact (ionization mode)
ELISA: Enzyme-Linked Immunosorbent Assay
EPM: Elevated Plus Maze
F1: First Filial Generation
FST: Forced Swim Test
GC/ECNI-MS: Gas Chromatography/Electron Capture Negative Ion Mass Spectrometry
gDNA: Genomic DNA
G × E: Gene–Environment Interaction
H-DE-71: DE-71 at 0.4mg/kg
HNS: Hypothalamic Neurohypophyseal System
HRGC/HRMS: High Resolution Gas Chromatography/High Resolution Mass Spectrometry
IQ: Intelligence Quotient
JWatcher: Java-Based Event Recording Software
LOQ: Limit of Quantification
L-DE-71: DE-71 at 0.1mg/kg
LS: Lateral Septum
l.w.: Lipid Weight
MB: Marble Burying
MDL: Method Detection Limit
MIQE: Minimum Information for Publication of Quantitative Real-Time PCR Experiments
MS: Mass Spectrometry
NDD: Neurodevelopmental Disorder
NOR: Novel Object Recognition
NTC: No-Template Control
OFT: Open Field Test
OD: Optical Density
OHT: Olfactory Habituation/Dishabituation Test
OPT: Olfactory Preference Test
OXT (*Oxt*): Oxytocin
*Oxtr*: Oxytocin Receptor
PBDE: Polybrominated Diphenyl Ether
PCR: Polymerase Chain Reaction
PND: Postnatal Day
POP: Persistent Organic Pollutant
ppm: Parts per Million
PVN: Paraventricular Nucleus of the hypothalamus
qPCR: Quantitative Polymerase Chain Reaction
RNA: Ribonucleic Acid
SEM / s.e.m.: Standard Error of the Mean
SMRT: Social Recognition Memory Test
SNP: Social Novelty Preference
SNN: Social neural network
SON: Supraoptic Nucleus
SRM: Social Recognition Memory
SRS: Social Reciprocity Score
SUOK: Elevated rod test for anxiety and sensorimotor behavior
T4: Thyroxine
TH: Thyroid Hormone
VEH/CON: Vehicle control
w.w.: Wet Weight

