## Supplementary Information for "Enduring Autism-like Phenotypes and Deregulated Hypothalamic Prosocial Peptides After Early-Life Exposure to Indoor Flame Retardants in Male C57BL/6 Mice"

**Supplementary Table 1**

GC/MS parameters for the isomer specific detection of PBDE.

| | |
|---|---|
| GC Type | Agilent 6890 |
| Column | Rtx-1614, 15 m, 0.25 mm ID, 0.1 µm film thickness (Restek) |
| Temperature program | 75 °C, 1.5 min, 18 °C/min, 210 °C, 8 °C/min, 310 °C, 5 min |
| Carrier gas | helium |
| Flow | constant flow 1.6 mL/min |
| Injector | cooled injection system CIS 4 (Gerstel) |
| Temperature transfer line | 320 °C |
| Injection volume | 1 µL splitless |
| MS Type | MAT 95XL (Thermo) |
| Ionization mode | EI+, 47 eV, 260°C |
| Resolution | >9000 |

**Supplementary Table 2**. Limits of quantification for PND 30 samples. **(Related to Figure 1)**

| | Mean | Standard deviation |
|---|---|---|
| | µg/kg wet weight | µg/kg wet weight |
| BDE-7 | 0.008 | 0.006 |
| BDE-10 | 0.006 | 0.005 |
| BDE-15 | 0.15 | 0.04 |
| BDE-17 | 0.008 | 0.002 |
| BDE-28 | 0.15 | 0.04 |
| BDE-30 | 0.01 | 0.01 |
| BDE-47 | 1.5 | 0.37 |
| BDE-49 | 0.02 | 0.006 |
| BDE-66 | 0.06 | 0.05 |
| BDE-71 | 0.02 | 0.004 |
| BDE-77 | 0.05 | 0.04 |
| BDE-85 | 0.05 | 0.04 |
| BDE-99 | 0.95 | 0.23 |
| BDE-100 | 0.43 | 0.10 |
| BDE-119 | 0.05 | 0.01 |
| BDE-126 | 0.05 | 0.04 |
| BDE-138 | 0.07 | 0.06 |
| BDE-139 | 0.06 | 0.05 |
| BDE-140 | 0.06 | 0.05 |
| BDE-153 | 0.75 | 0.18 |
| BDE-154 | 0.05 | 0.04 |
| BDE-156 | 0.07 | 0.06 |
| BDE-171 | 0.12 | 0.05 |
| BDE-180 | 0.12 | 0.05 |
| BDE-183 | 0.09 | 0.04 |
| BDE-184 | 0.09 | 0.04 |
| BDE-191 | 0.09 | 0.04 |
| BDE-196 | 0.02 | 0.02 |
| BDE-197 | 0.02 | 0.02 |
| BDE-201 | 0.02 | 0.02 |
| BDE-203 | 0.02 | 0.02 |
| BDE-204 | 0.02 | 0.02 |
| BDE-205 | 0.03 | 0.04 |
| BDE-206 | 0.31 | 0.07 |
| BDE-207 | 0.36 | 0.09 |
| BDE-208 | 0.21 | 0.05 |
| BDE-209 | 5.0 | 1.2 |

**Supplementary Table 3.** Mass spectrometric analysis (HRGC/HRMS) of PBDE congeners in PND 30 F1 male offspring brain on a wet-weight basis after transplacental and lactational low dose exposure to DE-71 through the dam. **(Related to Figure 1)**

| **Compound/Substituents** | **IUPAC Number** | **Male Offspring Brain PND 30 (ng/g ww)** | | |
|---|---|---|---|---|
| Treatment | | **VEH/CON** | **0.1 mg/kg DE-71** | **0.4 mg/kg DE-71** |
| n | | 3 | 3 | 3 |
| 2,4-Dibromdiphenylether | BDE-7 | <LOQ | <LOQ | NR |
| 2,6-Dibromdiphenylether | BDE-10 | <LOQ | <LOQ | <LOQ |
| 4,4'-Dibromdiphenylether | BDE-15 | <LOQ | <LOQ | <LOQ |
| 2,2',4-Tribromdiphenylether | BDE-17 | <LOQ | <LOQ | 0.015±0.003 |
| 2,4,4'-Tribromdiphenylether | BDE-28 | <LOQ | NR | <LOQ |
| 2,4,6-Tribromdiphenylether | BDE-30 | <LOQ | <LOQ | <LOQ |
| 2,2',4,4'-Tetrabromdiphenylether | BDE-47 | <LOQ | 5.625±4.014 | 8.344±2.125* |
| 2,2',4,5'-Tetrabromdiphenylether | BDE-49 | <LOQ | <LOQ | <LOQ |
| 2,3',4,4'-Tetrabromdiphenylether | BDE-66 | NR | <LOQ | <LOQ |
| 2,3',4',6-Tetrabromdiphenylether | BDE-71 | <LOQ | <LOQ | <LOQ |
| 3,3',4,4'-Tetrabromdiphenylether | BDE-77 | <LOQ | <LOQ | <LOQ |
| 2,2',3,4,4'-Pentabromdiphenylether | BDE-85 | 0.010±0.0003 | 1.014±0.644 | 1.238±0.087** |
| 2,2',4,4',5-Pentabromdiphenylether | BDE-99 | <LOQ | 16.400±9.083 | 46.843.5±7.706* |
| 2,2',4,4',6-Pentabromdiphenylether | BDE-100 | <LOQ | 6.007±3.292 | 20.065±3.30** |
| 2,3',4,4',6-Pentabromdiphenylether | BDE-119 | <LOQ | <LOQ | <LOQ |
| 3,3',4,4',5-Pentabromdiphenylether | BDE-126 | <LOQ | NR | <LOQ |
| 2,2',3,4,4',5'-Hexabromdiphenylether | BDE-138 | <LOQ | 0.358±0.235 | 0.612±0.117* |
| 2,2',3,4,4',6-Hexabromdiphenylether | BDE-139 | NR | 1.706±0.806 (*P*=.09) | 4.704±0.877* |
| 2,2',3,4,4',6'-Hexabromdiphenylether | BDE-140 | <LOQ | 0.391±0.327 | 0.541±0.132* |
| 2,2',4,4',5,5'-Hexabromdiphenylether | BDE-153 | <LOQ | 15.032±4.805* | 60.736±11.3* |
| 2,2',4,4',5,6'-Hexabromdiphenylether | BDE-154 | 0.018±0.001 | 1.214±0.666 | 2.801±0.493* |
| 2,3,3',4,4',5-Hexabromdiphenylether | BDE-156 | <LOQ | <LOQ | <LOQ |
| 2,2',3,3',4,4',6-Heptabromdiphenylether | BDE-171 | <LOQ | <LOQ | <LOQ |
| 2,2',3,4,4',5,5'-Heptabromdiphenylether | BDE-180 | <LOQ | <LOQ | <LOQ |
| 2,2',3,4,4',5',6-Heptabromdiphenylether | BDE-183 | <LOQ | NR | 0.214±0.008**** |
| 2,2',3,4,4',6,6'-Heptabromdiphenylether | BDE-184 | <LOQ | 0.119±0.069 | 0.353±0.09* |
| 2,3,3',4,4',5',6-Heptabromdiphenylether | BDE-191 | <LOQ | <LOQ | <LOQ |
| 2,2',3,3',4,4',5,6'-Octabromdiphenylether | BDE-196 | NR | 0.026±0.022 | <LOQ |
| 2,2',3,3',4,4',6,6'-Octabromdiphenylether | BDE-197 | NR | 0.038±0.034 | 0.034±0.014 |
| 2,2',3,3',4,5',6,6'-Octabromdiphenylether | BDE-201 | <LOQ | NR | <LOQ |
| 2,2',3,4,4',5',5',6-Octabromdiphenylether | BDE-203 | NR | 0.032±0.027 | <LOQ |
| 2,2',3,4,4',5',6,6'-Octabromdiphenylether | BDE-204 | <LOQ | NR | <LOQ |
| 2,3,3',4,4',5,5',6-Octabromdiphenylether | BDE-205 | <LOQ | <LOQ | <LOQ |
| 2,2',3,3',4,4',5,5',6-Nonabromdiphenylether | BDE-206 | <LOQ | <LOQ | <LOQ |
| 2,2',3,3',4,4',5,6,6'-Nonabromdiphenylether | BDE-207 | <LOQ | <LOQ | <LOQ |
| 2,2',3,3',4,5,5',6,6'-Nonabromdiphenylether | BDE-208 | <LOQ | <LOQ | <LOQ |
| 2,2',3,3',4,4',5,5',6,6'-Decabromdiphenylether | BDE-209 | <LOQ | <LOQ | <LOQ |
| ΣPBDEs | | 0.038±0.009 | 45.4±21.0* | 147±26.1**,^ |

Content was measured as ng/g wet weight
Values expressed as mean ± S.E.M.

Values for all biological replicates are listed; all values were identical when <LOQ is listed
**P*<.05, ***P*<.01 compared to VEH/CON; ^*P*<.05 compared to L-DE-71 (Student's t-test with Welch's Correction)
'n' indicates the number of biological replicates per group.
BDE, brominated diphenyl ether; IUPAC, International Union of Pure and Applied Chemistry; LOQ, limit of quantification; NR, not repeatable, only 1 sample yielded detectable values

**Supplementary Table 4.** Mass spectrometric analysis (GC/ECNI/MS) of PBDE congeners in PND 110 F1 male offspring brain on a lipid-weight basis after transplacental and lactational low dose exposure to DE-71 through the dam. **(Related to Figure 1)**

| Compound/Substituents | IUPAC Number | Male Offspring Brain PND 110 (ng/g lw) | | |
|---|---|---|---|---|
| **Treatment** | | **VEH/CON** | **0.1 mg/kg DE-71** | **0.4 mg/kg DE-71** |
| n | | 4 | 4 | 4 |
| 2,2′,4-tri BDE | BDE 17 | <MDL | <MDL | <MDL |
| 2,3′,4-tri BDE | BDE 25 | <MDL | <MDL | <MDL |
| 2,4,4′-tri BDE, 2′,3,4-tri BDE | BDE 28, 33 | <MDL | 13.27, 8.66, 10.14, 18.02 (12.5 ± 2.01)** | <MDL |
| 2,4,6,-tri BDE | BDE 30 | <MDL | | |
| 2,2′,4,4′-tetra BDE | BDE 47 | <MDL | 29.41, 23.29, 15.04, 40.78 (27.1 ± 5.42)** | 42.07, 106.31, 57.02, 47.97 (63.3 ± 14.6)* |
| 2,2′,4,5′-tetra BDE | BDE 49 | <MDL | <MDL | <MDL |
| 2,3′,4,4′-tetra BDE | BDE 66 | <MDL | <MDL | <MDL |
| 2,3′,4′,6-tetra BDE | BDE 71 | <MDL | <MDL | <MDL |
| 2,4,4′,6-tetra BDE | BDE 75 | <MDL | <MDL | <MDL |
| 2,2′,3,4,4′-penta BDE, 2,2′,4,4′,6,6′-hexa BDE | BDE 85, 155 | <MDL | <MDL | <MDL |
| 2,2′,4,4′,5-penta BDE | BDE 99 | <MDL | #55.59, <MDL, <MDL, <MDL | <MDL |
| 2,2′,4,4′,6-penta BDE | BDE 100 | <MDL | 14.70, 5.96. 5.56, 67.33 (23.4 ± 14.8) | 10.76, 27.20, 10.61, 13.71 (15.6 ± 3.94)* |
| 2,3,4,5,6-penta BDE | BDE 116 | <MDL | <MDL | <MDL |
| 2,3′,4,4′,6 –penta BDE | BDE 119 | <MDL | <MDL | <MDL |
| 2,2′,3,4,4′,5′-hexa BDE | BDE 138 | <MDL | <MDL | <MDL |
| 2,2′,4,4′,5,5′-hexa BDE (mean ± S.E.M.) | BDE 153 | <MDL | 10.76, 11.37, 5.89, 34.14 (15.5 ± 6.32)* | 129.14, 222.51, 135.25, 117.49 (151 ± 24.1)** |
| 2,2′,4,4′,5,6′-hexa BDE | BDE 154 | <MDL | <MDL | <MDL |
| 2, 3,3′,4,4′,5-hexa BDE | BDE 156 | <MDL | <MDL | <MDL |
| 2,2′, 3,4,4′,5,6-hepta BDE | BDE 181 | <MDL | <MDL | <MDL |
| 2,2′, 3, 4,4′,5′,6-hepta BDE | BDE 183 | <MDL | <MDL | <MDL |
| 2,3,3′,4,4′,5,6-hepta BDE | BDE 190 | <MDL | <MDL | <MDL |
| 2, 3,3′,4,4′,5′,6-hepta BDE | BDE 191 | <MDL | <MDL | <MDL |
| 2,2′,3,3′,4,5,6,6′-octa BDE, 2,2′,3,4,4′,5,5′,6-octa BDE | BDE 200, 203 | <MDL | <MDL | <MDL |
| 2, 3,3′,4,4′,5,5′,6-octa BDE | BDE 205 | <MDL | <MDL | <MDL |
| 2,2′,3,3′,4,4′,5,5′,6-nona BDE | BDE 206 | <MDL | <MDL | <MDL |
| 2,2′,3,3′,4,4′,5,5′,6,6′-deca BDE | BDE 209 | <MDL | <MDL | <MDL |
| **ΣPBDEs** | | <MDL | 78.6 ± 28.0* | 230 ± 42.3**,^ |

Content was measured as ng/g lipid weight
Values expressed as mean ± S.E.M.
Values for all biological replicates are listed; all values were identical when <MDL is listed
*$P$<.05, **$P$<.01 compared to VEH/CON (Student’s t-test with Welch’s Correction)
^$P$<.05 compared to L-DE-71 (Student’s t-test with Welch’s Correction)
‘n’ indicates the number of biological replicates per group. BDE-99 was detected in only one 0.1 mg/kg DE-71 biological replicate.
BDE, brominated diphenyl ether; IUPAC, International Union of Pure and Applied Chemistry; MDL, method detection limit

**Supplementary Table 5.** Mass spectrometric analysis (GC/ECNI/MS) of PBDE congeners in PND 110 F1 male offspring brain on a wet-weight basis after transplacental and lactational low dose exposure to DE-71 through the dam. **(Related to Figure 1)**

| Compound/Substituents | IUPAC Number | Male Offspring Brain PND 110 (ng/g ww) | | |
|---|---|---|---|---|
| Treatment | | VEH/CON | 0.1 mg/kg DE-71 | 0.4 mg/kg DE-71 |
| n | | 4 | 4 | 4 |
| 2,2′,4-tri BDE | BDE 17 | <MDL | <MDL | <MDL |
| 2,3′,4-tri BDE | BDE 25 | <MDL | <MDL | <MDL |
| 2,4,4′-tri BDE,<br>2′,3,4-tri BDE | BDE 28, 33 | <MDL | 0.30, 0.13, 0.25, 0.17<br>(0.213 ± 0.038)** | <MDL |
| 2,4,6,-tri BDE | BDE 30 | <MDL | <MDL | <MDL |
| 2,2′,4,4′-tetra BDE | BDE 47 | <MDL | 0.67, 0.35, 0.37, 0.38<br>(0.443 ± 0.076)** | 0.57, 0.56, 0.56, 0.55<br>(0.560 ± 0.004)**** |
| 2,2′,4,5′-tetra BDE | BDE 49 | <MDL | <MDL | <MDL |
| 2,3′,4,4′-tetra BDE | BDE 66 | <MDL | <MDL | <MDL |
| 2,3′,4′,6-tetra BDE | BDE 71 | <MDL | <MDL | <MDL |
| 2,4,4′,6-tetra BDE | BDE 75 | <MDL | <MDL | <MDL |
| 2,2′,3,4,4′-penta BDE,<br>2,2′,4,4′,6,6′-hexa BDE | BDE 85, 155 | <MDL | <MDL | <MDL |
| 2,2′,4,4′,5-penta BDE | BDE 99 | <MDL | #1.27, <MDL, <MDL,<br><MDL | <MDL |
| 2,2′,4,4′,6-penta BDE | BDE 100 | <MDL | 0.34, 0.09, 0.14, 0.62<br>(0.298 ± 0.12) *P*=.06 | 0.15, 0.14, 0.10, 0.16<br>(0.138 ± 0.013)** |
| 2,3,4,5,6-penta BDE | BDE 116 | <MDL | <MDL | <MDL |
| 2,3′,4,4′,6 –penta BDE | BDE 119 | <MDL | <MDL | <MDL |
| 2,2′,3,4,4′,5′-hexa BDE | BDE 138 | <MDL | <MDL | <MDL |
| 2,2′,4,4′,5,5′-hexa BDE<br>(mean ± S.E.M.) | BDE 153 | <MDL | 0.25, 0.17, 0.14, 0.31<br>(0.218 ± 0.039)** | 1.76, 1.16, 1.32, 1.36<br>(1.40 ± 0.128)*** |
| 2,2′,4,4′,5,6′-hexa BDE | BDE 154 | <MDL | <MDL | <MDL |
| 2, 3,3′,4,4′,5-hexa BDE | BDE 156 | <MDL | <MDL | <MDL |
| 2,2′, 3,4,4′,5,6-hepta BDE | BDE 181 | <MDL | <MDL | <MDL |
| 2,2′, 3, 4,4′,5′,6-hepta BDE | BDE 183 | <MDL | <MDL | <MDL |
| 2,3,3′,4,4′,5,6-hepta BDE | BDE 190 | <MDL | <MDL | <MDL |
| 2, 3,3′,4,4′,5′,6-hepta BDE | BDE 191 | <MDL | <MDL | <MDL |
| 2,2′,3,3′,4,5,6,6′-octa BDE,<br>2,2′,3,4,4′,5,5′,6-octa BDE | BDE 200, 203 | <MDL | <MDL | <MDL |
| 2, 3,3′,4,4′,5,5′,6-octa BDE | BDE 205 | <MDL | <MDL | <MDL |
| 2,2′,3,3′,4,4′,5,5′,6-nona BDE | BDE 206 | <MDL | <MDL | <MDL |
| 2,2′,3,3′,4,4′,5,5′,6,6′-deca BDE | BDE 209 | <MDL | <MDL | <MDL |
| ΣPBDEs | | <MDL | 1.17 ± 0.205* | 2.10 ± 0.134**,^ |

Content was measured as ng/g wet weight

Values expressed as mean ± S.E.M.

Values for all biological replicates are listed; all values were identical when <MDL is listed

**P*<.05, ***P*<.01, ****P*<.001, *****P*<.0001 compared to VEH/CON (Student's t-test with or without Welch's Correction or Brown-Forsythe ANOVA)

^*P*<.05 compared to L-DE-71 (Student's t-test with Welch's Correction)

'n' indicates the number of biological replicates per group. BDE-99 was detected in only one 0.1 mg/kg DE-71 replicate.

BDE, brominated diphenyl ether; IUPAC, International Union of Pure and Applied Chemistry; MDL, method detection limit

**Supplementary Figure 1**

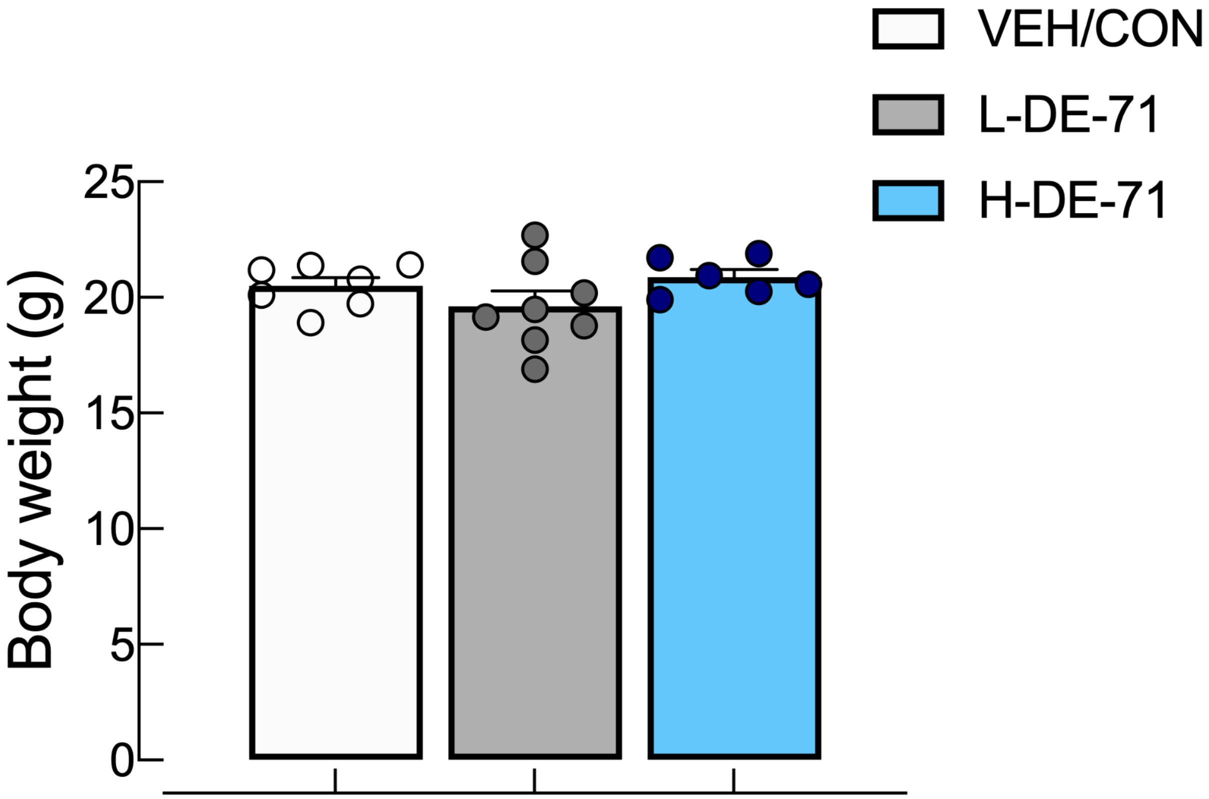


**Supplementary Figure 1,** Average body weights (mean +/- s.e.m) of male offspring at PND 44. *n*, 7-8 litters/group

**Supplementary Figure 2**

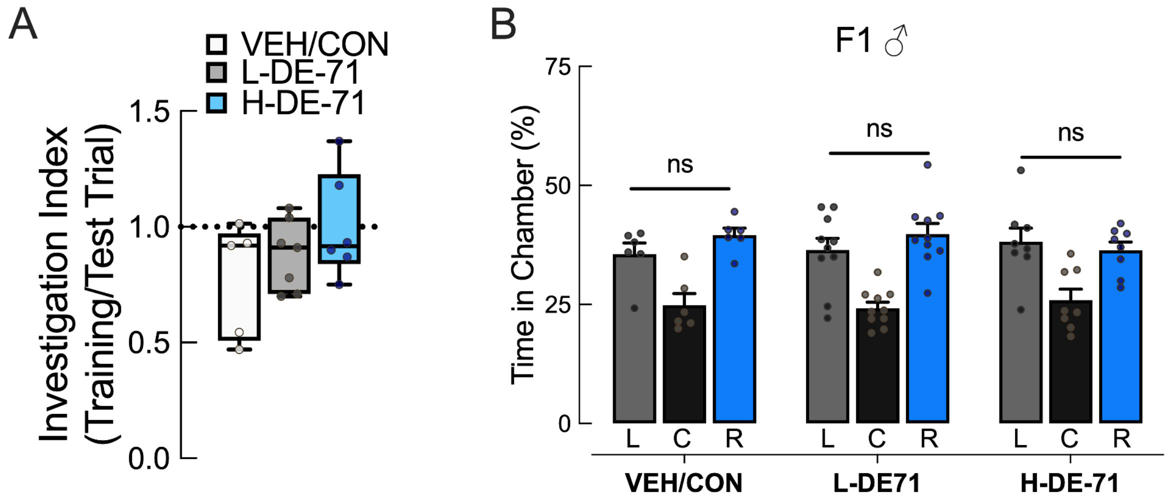


**Supplementary Figure 2.** Social novelty preference test (SNP) scores are not confounded by lack of sociability or sidedness on the three-chamber sociability test (SOC). **(**Related to Figure 2) **a** Investigation Index scores on SNP (when exploring both social stimuli) indicate the ratio of time spent investigating the first social stimulus on training trial to the total investigation time during test trial. A score of near 1 shown here indicates equal investigation time on training and test trials indicating that SNP deficit in DE-71 mice is not due to lack of social engagement. **b** Adult male mice were tested for sociability using a 3-chamber apparatus. In the first habituation phase, mice are allowed to explore the middle chamber only. During the second habituation phase a test mouse is allowed to explore all three chambers of an empty apparatus and time spent in left, right and center chambers is recorded over 10 min. Males in all exposure groups showed similar times spent in left and right chambers during the second habituation phase indicating no inherent side preference. *n*, 5-7 litters/group (a), 6-9 litters/group (b). C, Center; L, Left; R, right

**Supplementary Figure 3**

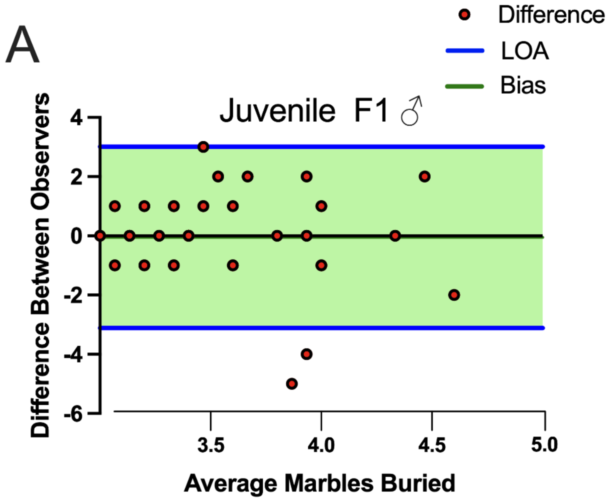


**Supplementary Figure 3.** Scores for marble burying and nestle shredding tests (Related to Figure 2) **a** Bland-Altman bias plot (mean+s.d.) was used to test the validity and reproducibility of marble burying scores between two independent judges blind to exposure group. Analysis revealed a very small mean of the differences between judge scores (Bias, -0.05±1.56) and a precision measured as limits of agreement (LOA), average difference ± 1.96 standard deviation of the difference, of -3.1-3.0, indicating negligible skewing by either judge. **b** Bland-Altman plot of adult marble burying (Bias, 0.37±3.51) LOA range from -6.5-7.2.

**Supplementary Figure 4**

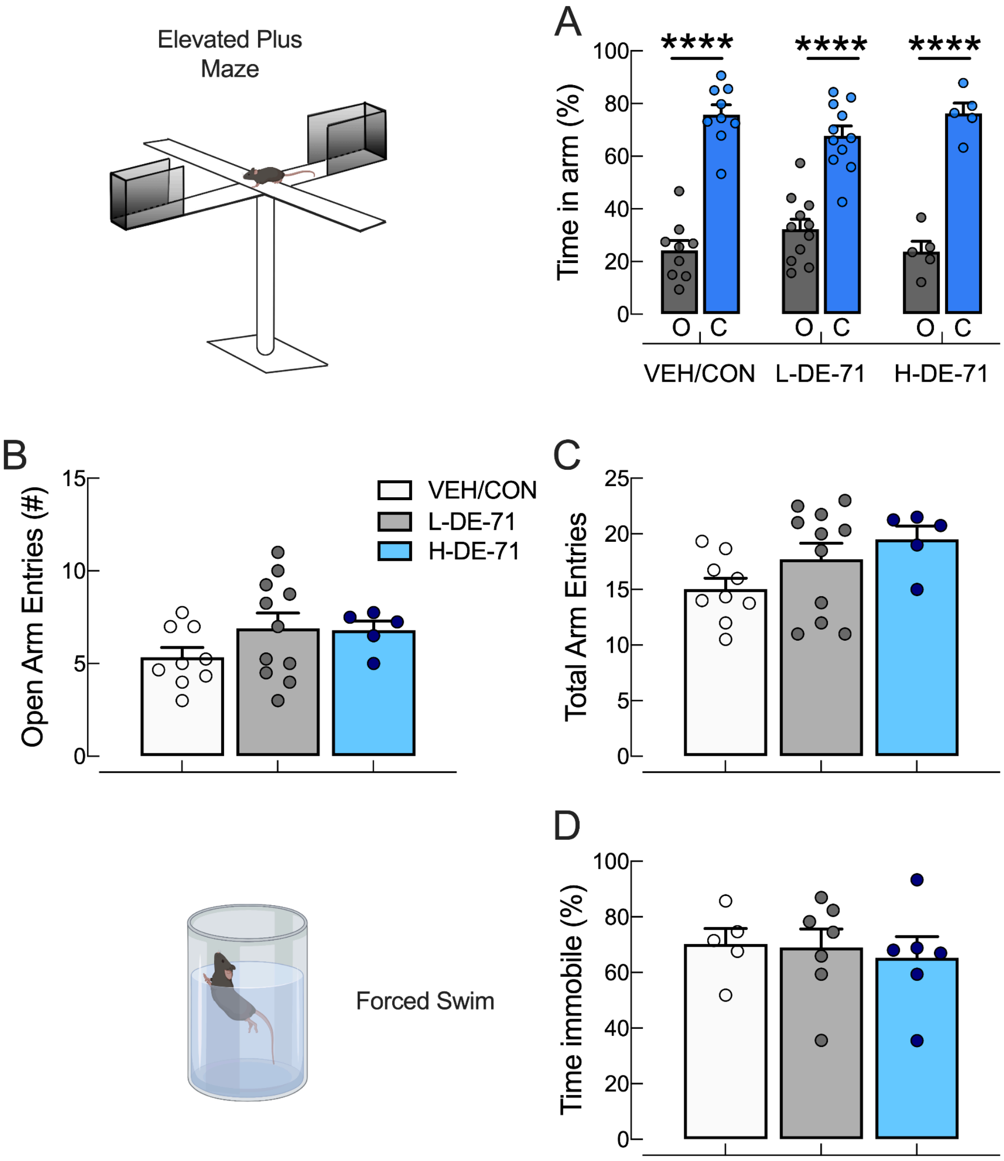


**Supplementary Figure 4. DE-71 exposure does not affect anxiety nor produce depressive-like behavior in exposed male offspring. a** Time spent in open and closed arms on an elevated plus maze. All groups, regardless of exposure, spent significantly greater time in closed arm. **b, c** Open arm and total arm entries were similar across all groups. **d** Time spent immobile on the Forced Swim test showed no differences across groups indicating no effects of DE-71 on depressive-like behavior. *compared to % time spent in closed arm, ****$P$<.0001. $n$, 5-11 litters/group (a, b, c); $n$, 5-7 litters/group (d).

**Supplementary Figure 5**

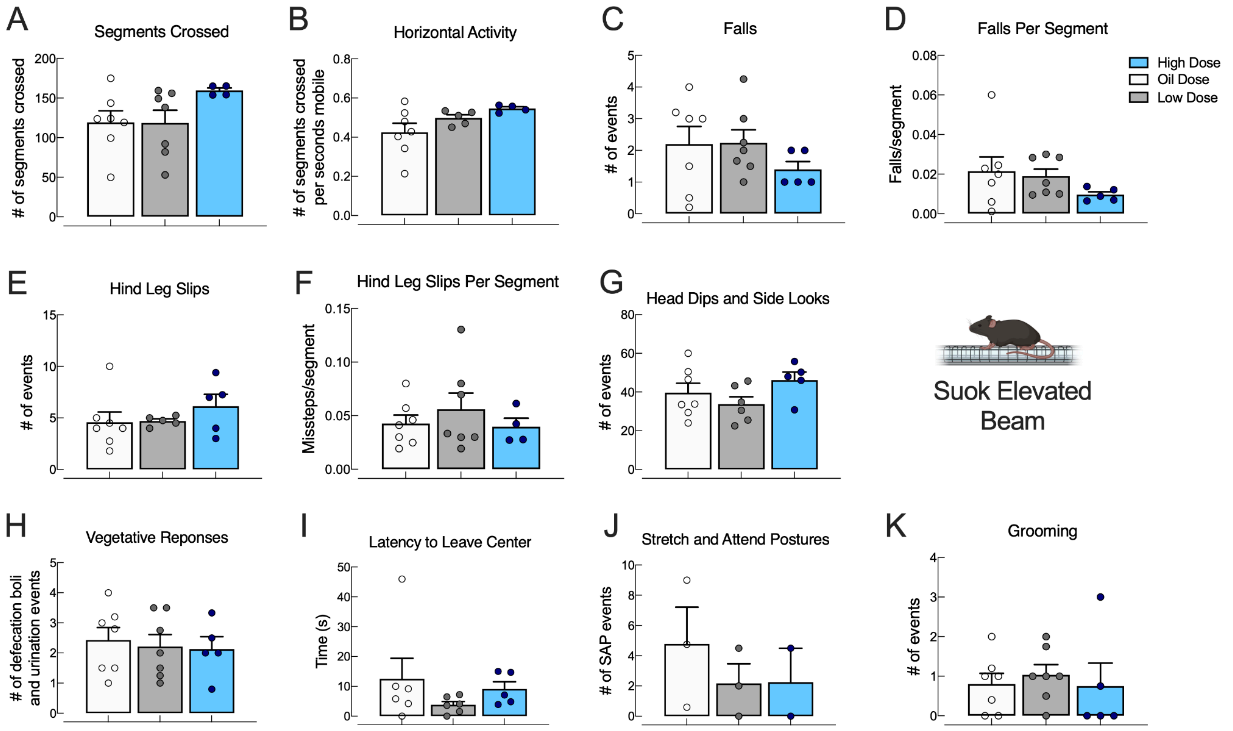


**Supplementary Figure 5** Suok Test Parameters. Male adults were tested on SUOK platform for different sensorimotor abilities: **a, b** locomotion; **c-f** sensorimotor coordination; **g** exploratory activity; **h-j** anxiety behaviors; and **k** autogrooming. *n* 6-8 litters/group (a-i, k); 2-3 litters/group (j)
